# Two transmembrane transcriptional regulators coordinate to activate chitin-induced natural transformation in *Vibrio cholerae*

**DOI:** 10.1101/2024.09.30.615920

**Authors:** Allison C. Hullinger, Virginia E. Green, Catherine A. Klancher, Triana N. Dalia, Ankur B. Dalia

## Abstract

Transcriptional regulators are a broad class of proteins that alter gene expression in response to environmental stimuli. Transmembrane transcriptional regulators (TTRs) are a subset of transcriptional regulators in bacteria that can directly regulate gene expression while remaining anchored in the membrane. Whether this constraint impacts the ability of TTRs to bind their DNA targets remains unclear. *Vibrio cholerae* uses two TTRs, ChiS and TfoS, to activate horizontal gene transfer by natural transformation in response to chitin by inducing the *tfoR* promoter (P*_tfoR_*). While TfoS was previously shown to bind and regulate P*_tfoR_* directly, the role of ChiS in P*_tfoR_* activation remains unclear. Here, we show that ChiS directly binds P*_tfoR_* upstream of TfoS, and that ChiS directly interacts with TfoS. By independently disrupting ChiS-P*_tfoR_* and ChiS-TfoS interactions, we show that ChiS-P*_tfoR_* interactions play the dominant role in P*_tfoR_* activation. Correspondingly, we show that in the absence of ChiS, recruitment of the P*_tfoR_* locus to the membrane is sufficient for P*_tfoR_* activation when TfoS is expressed at native levels. Finally, we show that the overexpression of TfoS can bypass the need for ChiS for P*_tfoR_* activation. All together, these data suggest a model whereby ChiS both (1) recruits the P*_tfoR_* DNA locus to the membrane for TfoS and (2) directly interacts with TfoS, thereby recruiting it to the membrane-proximal promoter. This work furthers our understanding of the molecular mechanisms that drive chitin-induced responses in *V. cholerae* and more broadly highlights how the membrane-embedded localization of TTRs can impact their activity.

**AUTHOR SUMMARY:** Living organisms inhabit diverse environments where they encounter a wide range of stressors. To survive, they must rapidly sense and respond to their surroundings. One universally conserved mechanism to respond to stimuli is via the action of DNA-binding transcriptional regulators. In bacterial species, these regulators are canonically cytoplasmic proteins that freely diffuse within the cytoplasm. In contrast, an emerging class of transmembrane transcriptional regulators (TTRs) directly regulate gene expression from the cell membrane. Prior work shows that two TTRs, TfoS and ChiS, cooperate to activate horizontal gene transfer by natural transformation in response to chitin in the facultative pathogen *Vibrio cholerae*. However, how these TTRs coordinate to activate this response has remained unclear. Here, we show that ChiS likely promotes TfoS-dependent activation of natural transformation by (1) relocalizing its target promoter to the membrane and (2) recruiting TfoS to the membrane proximal promoter through a direct interaction. Together, these results inform our understanding of both the *V. cholerae* chitin response and how TTR function can be impacted by their membrane localization.

## INTRODUCTION

Bacterial species inhabit diverse environments where they encounter a wide range of stressors. To survive fluctuating conditions, they must have ways to sense and respond to environmental change. One such mechanism is the alteration of gene expression in response to specific environmental cues using a broad group of proteins called transcriptional regulators. Most commonly, transcriptional regulators are localized to the cytoplasm, where they can freely diffuse to find their DNA targets. However, recent work highlights the presence of transmembrane transcriptional regulators (TTRs) in diverse bacterial genomes [1]. In the few examples where they have been studied, TTRs were shown to bind to their DNA targets while remaining anchored in the membrane [2–7]. However, TTRs remain relatively poorly characterized, and the factors that influence how these proteins find their DNA targets to regulate gene expression remain unclear. Here, we investigate the mechanism of action for two chitin-responsive TTRs in *Vibrio cholerae*—TfoS and ChiS.

Natural transformation is a broadly conserved mode of horizontal gene transfer in which cells take up exogenous DNA to then integrate into their genome by homologous recombination. This process facilitates the rapid transfer of beneficial traits in diverse bacterial species. TfoS is a TTR that is required for the induction of natural transformation in *V. cholerae*. In addition to TfoS, induction of natural transformation requires chitin oligosaccharides, which *V. cholerae* liberates from the shells of crustacean zooplankton in the marine environment [8, 9]. Previous work shows that in the presence of chitin oligosaccharides, TfoS directly activates the expression of the small RNA (sRNA) *tfoR* by binding to its promoter (P*_tfoR_*) [8, 9]. When transcribed, *tfoR* base pairs with the 5’ untranslated region (UTR) of the mRNA of the master regulator of competence, TfoX, which allows for its translation [10]. TfoX then activates the expression of the genes necessary for DNA uptake and integration [11, 12]. Although this process requires chitin, it remains unclear whether TfoS senses chitin directly or indirectly through an intermediate.

ChiS is a TTR that is critical for induction of the chitin utilization program, a cascade of genes required for chitin uptake and catabolism that facilitates *V. cholerae* survival in marine habitats [13, 14]. When soluble chitin oligosaccharides enter the periplasm, they are bound by chitin-binding protein (CBP) [14, 15]. Chitin-bound CBP then activates ChiS, likely via a direct interaction [4]. Activated ChiS can then bind to the promoter of the chitobiose (*chb*) utilization operon (P*_chb_*) to activate the expression of genes required for chitin uptake and catabolism [4, 14]. Like TfoS, ChiS is necessary for chitin-induced natural transformation [8, 9]. However, it is unclear whether this is due to its role in promoting chitin catabolism or whether it more directly regulates the genes required for natural transformation. Previous studies demonstrate that ChiS and TfoS are both required for chitin-induced P*_tfoR_* activation, supporting the latter possibility [8, 9]. However, the mechanism by which ChiS and TfoS coordinate to regulate P*_tfoR_* expression remains unclear.

## RESULTS

### ChiS binds to P_tfoR_ and is required for P_tfoR_ activation

Prior work suggests that ChiS and TfoS are both required for activation of P*_tfoR_* [8, 9]. To formally test this, we generated strains with a chromosomally integrated P*_tfoR_*-*gfp* transcriptional reporter. These strains also contained a previously described P*_chb_*-*mCherry* construct [16], which serves as a reporter for ChiS activity. In addition, these strains contained a constitutively expressed [17] mTFP1 construct (P*_const2_*-*mTFP1*), which was used to normalize for intrinsic noise in gene expression. All reporter constructs were chromosomally integrated at ectopic genomic sites. Because ChiS and TfoS are TTRs, the genomic locus where these transcriptional reporters are integrated could influence their activation (*i.e.*, they could be poorly activated if inserted into regions of the genome that are inaccessible to TTRs). It was therefore crucial to verify that these reporters could still be activated by their respective TTRs. When a strain harboring these constructs was incubated on chitin, we observed strong activation of both P*_tfoR_* and P*_chb_* by fluorescence microscopy (**Fig. 1A-C**). This result is consistent with their known regulation by chitin [4, 10], suggesting that these chromosomally integrated reporters are functional. Deleting either *chiS* or *tfoS* resulted in a complete loss of P*_tfoR_* activation (**Fig. 1A-B**). Furthermore, we found that P*_chb_* expression was completely ablated by Δ*chiS*, but not by Δ*tfoS* (**Fig. 1C**), which is consistent with the previously established role for ChiS in P*_chb_* activation [4, 14]. These results confirm that both ChiS and TfoS are required for P*_tfoR_* induction and demonstrate that these TTRs are not equally required for expression of all chitin-regulated promoters in *V. cholerae*. Furthermore, these results verify that the ectopic P*_tfoR_* and P*_chb_* reporter constructs can be activated by these TTRs.

**Fig. 1.**
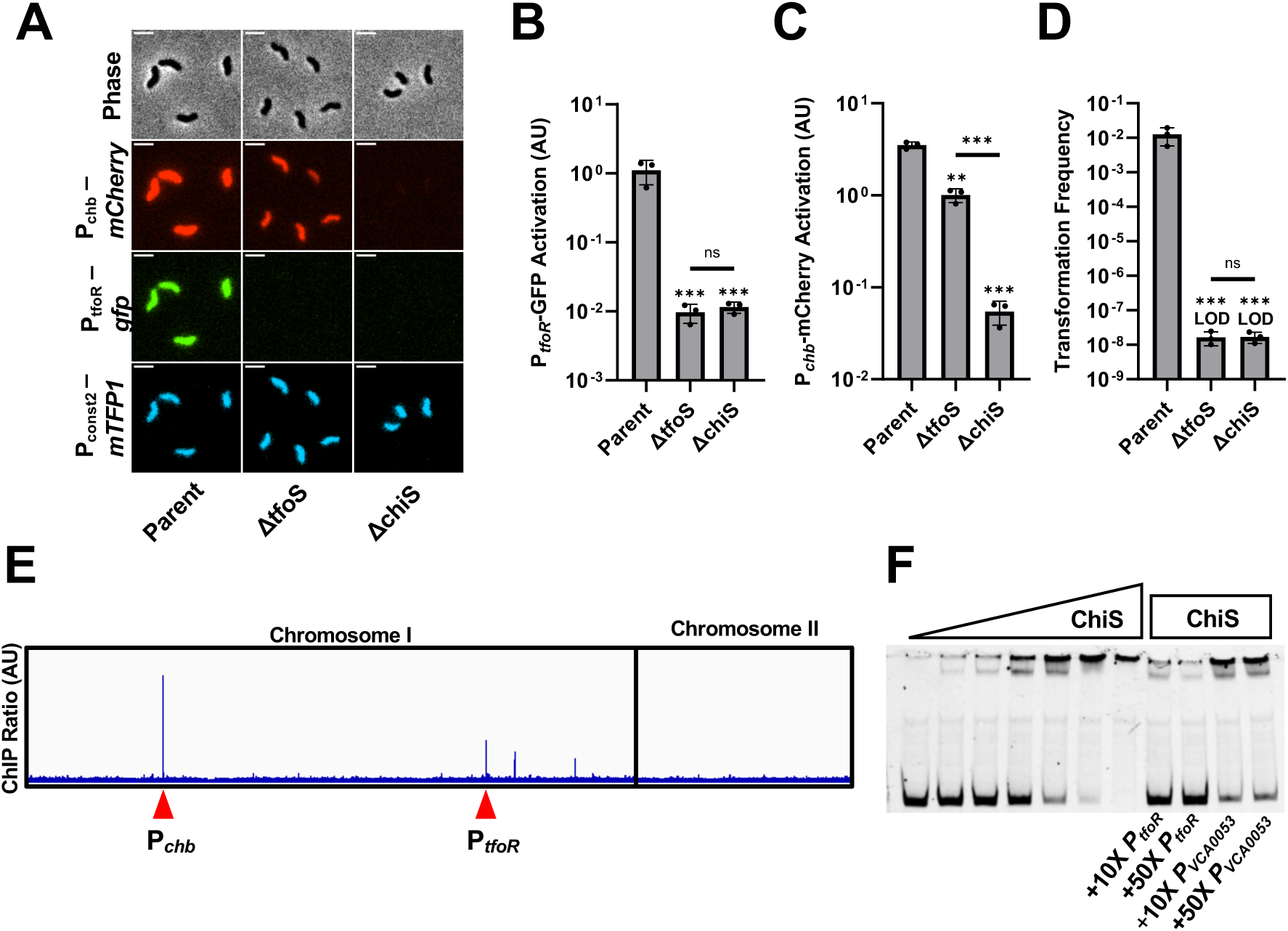
ChiS binds P_tfoR_ and is necessary for P_tfoR_ activation. (**A-C**) Transcriptional reporter assays to assess chitin-dependent gene expression. The indicated strains all harbored 3 fluorescent reporter constructs: P*_chb_-mCherry*, P*_tfoR_-gfp*, and P*_const2_-mTFP1*. Cells were incubated on chitin and then imaged via epifluorescence microscopy to assess reporter expression. Representative images are shown in **A** and the quantification of the results are shown in **B** and **C**. Scale bar, 2µm. For each replicate (*n* = 3), the geometric mean fluorescence was determined by analyzing 300 individual cells. (**D**) Chitin-induced natural transformation assay of the indicated strains. Results are from three independent biological replicates and shown as the mean ± SD. Statistical comparisons in **B**, **C**, and **D** were made by one-way ANOVA with Tukey’s multiple comparison test on log-transformed data (normal distribution confirmed by Shapiro-Wilk test). Statistical identifiers directly above bars represent comparisons to the parent. ns, not significant. *** = *p* < 0.001, ** = *p* < 0.01. LOD, limit of detection. (**E**) ChIP-seq analysis uncovers ChiS binding sites. The panel shows the “ChIP ratio”, which is defined as the mapping distribution of ChIP-enriched DNA normalized to the mapping distribution of input DNA. ChIP peaks at the P*_chb_* and P*_tfoR_* loci are denoted by a red arrow. (**F**) EMSAs to test ChiS binding at P*_tfoR_*. Increasing concentrations of the purified ChiS cytoplasmic domain (from left to right: 0 nM, 25 nM, 50 nM, 100 nM, 200 nM, 400 nM, 800 nM) were incubated with Cy5-labeled P*_tfoR_* DNA probe. Cold competitor EMSAs were carried out using 200 nM ChiS and either 10-fold or 50-fold excess of the unlabeled competitor DNA probe as indicated. Data in **F** are representative of three independent experiments.

The downstream consequence of P*_tfoR_* induction is the expression of the genes necessary for horizontal gene transfer by natural transformation. We therefore also assessed the phenotypic impact of P*_tfoR_* regulation by ChiS and TfoS on natural transformation. We found that deletion of either *chiS* or *tfoS* reduced natural transformation to the limit of detection (**Fig. 1D**), which is consistent with a critical role for both TTRs in inducing this behavior. Further, the overexpression of *tfoR*, which can induce competence in the absence of chitin [18–20], restores natural transformation in both Δ*chiS* and Δ*tfoS* backgrounds, suggesting that the inability of these strains to transform is due to the lack of *tfoR* expression (**Fig. S1**).

Our previous work has established that TfoS directly binds P*_tfoR_* and that the ectopic expression of TfoS in the heterologous host *E. coli* is sufficient to drive P*_tfoR_* induction even in the absence of ChiS [8]. However, our results above demonstrate that ChiS is still necessary for P*_tfoR_* activation in *V. cholerae*. Thus, one possibility is that ChiS is necessary to stabilize natively expressed TfoS in *V. cholerae* (*i.e.,* that native TfoS is degraded in the absence of ChiS). To test this hypothesis, we generated a functional FLAG-tagged allele of TfoS (**Fig. S2**) and performed Western blot analysis. We found that TfoS expression is not altered in the Δ*chiS* background (**Fig. S2C-D**). This result suggests that ChiS does not promote P*_tfoR_* activation by simply stabilizing TfoS.

Because ChiS does not stabilize TfoS, we reasoned that it may play a more active role in regulating P*_tfoR_*. Specifically, we hypothesized that ChiS may directly bind to P*_tfoR_*. To test this possibility, we performed chromatin immunoprecipitation-sequencing (ChIP-seq) to identify ChiS binding sites *in vivo*. Consistent with our previous demonstrating that ChiS binds to P*_chb_* [4], we observed a strong ChIP peak at the P*_chb_* promoter (**Fig. 1E**). ChIP peaks were also identified upstream of a predicted chitinase gene (VC2217) as well as a putative glycosyltransferase (VC2487). Strikingly, this analysis also revealed a strong ChIP peak at P*_tfoR_* (**Fig. 1E**). To verify this result, we performed electrophoretic mobility shift assays (EMSAs) using the purified cytoplasmic domain of ChiS and found that it binds to P*_tfoR_ in vitro* (**Fig. 1F**). Furthermore, we show that this interaction could be competitively inhibited by unlabeled P*_tfoR_* probe but not by a nonspecific promoter probe (P*_VCA0053_*) (**Fig. 1F**), indicating that the interaction of ChiS with P*_tfoR_* is specific. Together, these results indicate that both ChiS and TfoS bind P*_tfoR_*, and that both TTRs are critical for chitin-induced expression of TfoR.

### ChiS directly interacts with TfoS

Our results thus far are consistent with two possible models for activation of P*_tfoR_* by ChiS and TfoS. One possibility is that TfoS cannot bind P*_tfoR_* independently, but instead relies on ChiS binding at P*_tfoR_* to recruit this DNA locus to the membrane so that it becomes accessible to TfoS. Alternatively, ChiS may bind alongside TfoS at P*_tfoR_* and then directly interact with TfoS to allosterically regulate its activity by inducing a conformational change that allows TfoS to bind P*_tfoR_*. One of the primary differences between these two models is that the latter has a strict requirement for a direct interaction between ChiS and TfoS, while the former model does not. To test whether ChiS and TfoS interact, we performed PopZ linked apical recruitment (POLAR) assays [21]. In these assays, two putatively interacting proteins, the “bait” and the “prey”, are translationally fused to distinct fluorescent markers to track their independent localization. The bait protein is also fused to an H3H4 PopZ interaction domain derived from *Caulobacter crescentus*. Upon induction of the polarly localizing PopZ protein, the bait is relocalized to the cell pole. If the prey interacts with the bait, it will also relocalize to the cell pole. If there is no interaction between the bait and prey, the localization of the prey should be unaffected by PopZ induction. To assess interactions between ChiS and TfoS, we generated strains expressing ChiS-msfGFP-H3H4, TfoS-mCherry, and arabinose-inducible PopZ. In the absence of ChiS, TfoS-mCherry was diffusely localized at the cell periphery, consistent with it being a membrane-embedded regulator, and this localization did not change when PopZ was induced (**Fig. 2**). We have previously found that ChiS-msfGFP-H3H4 inherently forms foci at the cell periphery for reasons that remain unclear [4], and we have recapitulated this observation here (**Fig. 2**). We found that in cells expressing both ChiS-msfGFP-H3H4 and TfoS-mCherry, TfoS-mCherry forms foci that strongly colocalize with ChiS-msfGFP-H3H4, suggesting that these proteins may interact (**Fig. 2**). Furthermore, when PopZ is induced in these cells, both ChiS-msfGFP-H3H4 and TfoS-mCherry relocalize to the cell poles, which is consistent with ChiS-TfoS interactions (**Fig. 2**). As an independent way to assess ChiS-TfoS interactions, we also performed bacterial adenylate cyclase two-hybrid (BACTH) assays [22]. Briefly, this assay involves translationally fusing bait and prey proteins to each half of a split adenylate cyclase reporter. Physical interaction between the bait and prey results in reconstitution of adenylate cyclase activity, which induces beta-galactosidase expression in the *E. coli* reporter strain. These BACTH assays also showed an interaction between ChiS and TfoS, further suggesting that these proteins directly interact (**Fig. S3**).

**Fig. 2.**
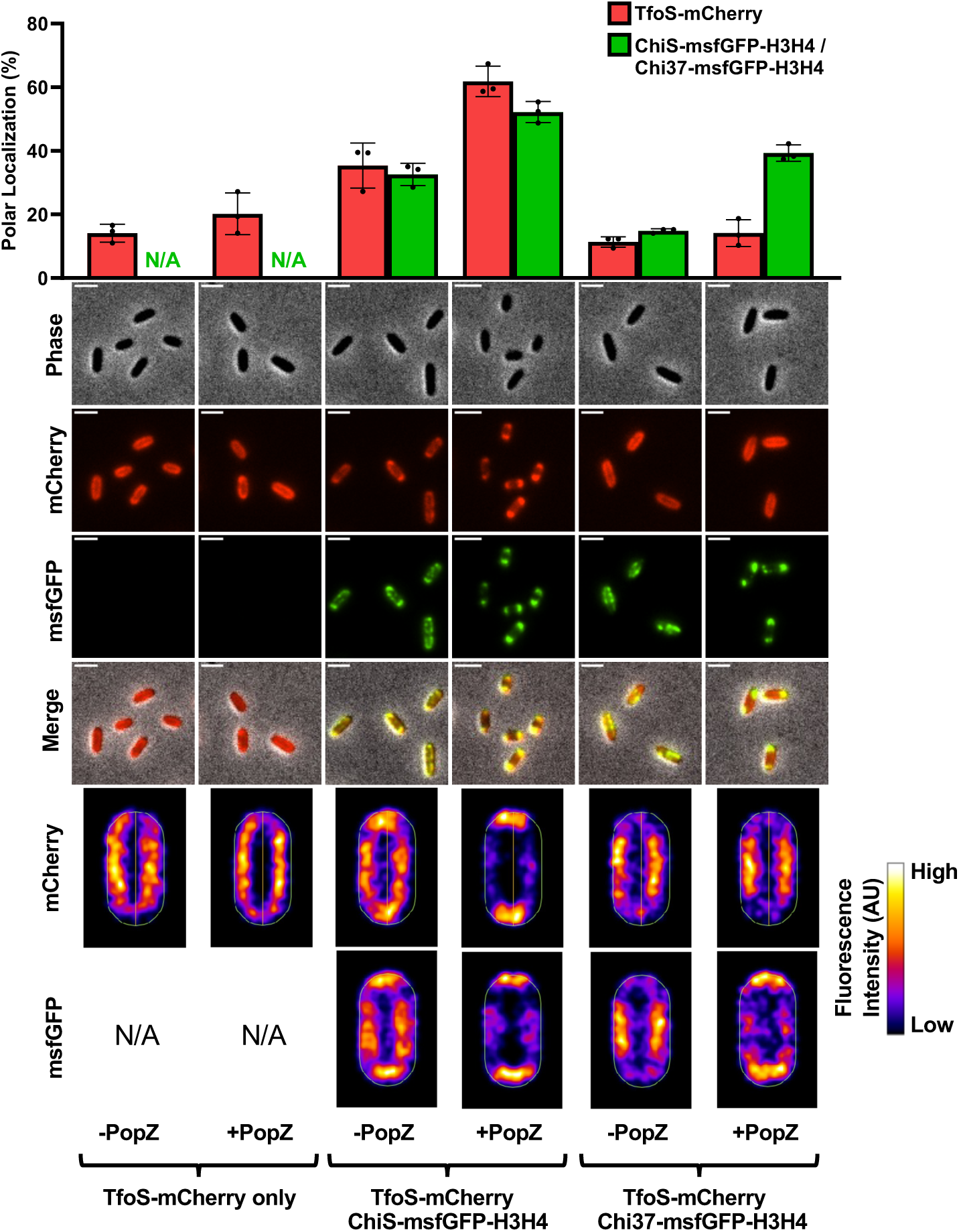
ChiS interacts with TfoS. POLAR assays were conducted to assess the interaction between ChiS and TfoS. Colocalization was assessed in cells containing P*_bad_-popZ* and the indicated ChiS and TfoS fusion constructs. The localization of these constructs was assessed in the absence (-PopZ) and presence (+PopZ) of PopZ induction. Scale bar, 2µm. Representative images of cells are shown along with heat maps that represent the fluorescence localization in 300 cells. Data are representative of three independent biological replicates. The proportion of TfoS-mCherry (red bars) and ChiS-msfGFP-H3H4 / Chi37-msfGFP-H3H4 (green bars) fluorescence within the polar region (≤ 0.2 µm from the cell pole) was quantified. For each replicate (*n* = 3), fluorescence localization was determined by analyzing 300 individual cells. N/A, not applicable

### ChiS-TfoS interactions support, but are not critical for P_tfoR_ activation

We next wanted to test whether the interaction between ChiS and TfoS is critical for P*_tfoR_* activation, so we sought to disrupt the interaction between these proteins. To that end, we created a functional chimera called Chi37, which is composed of the two transmembrane domains and a truncated portion of the periplasmic domain of the chemoreceptor Mlp37 translationally fused to the cytoplasmic domain of ChiS. Chi37 is functional for P*_chb_* activation and binds both P*_chb_* and P*_tfoR_* at wildtype levels *in vivo* according to ChIP-qPCR assays (**Fig. S4**). However, POLAR assays demonstrate that Chi37-msfGFP-H3H4 cannot relocalize TfoS-mCherry, indicating that Chi37-TfoS interactions are below the limit of detection of this assay (**Fig. 2**). Because Chi37 retains the ability to bind DNA but exhibits diminished interactions with TfoS, we used this allele to test the impact of ChiS-TfoS interactions on P*_tfoR_* activation.

If the interaction between ChiS and TfoS is essential for P*_tfoR_* activation, we hypothesized that a strain harboring Chi37 will not be able to activate P*_tfoR_* expression, and therefore will not allow for chitin-induced natural transformation. However, we found that a Chi37 mutant had an intermediate phenotype – it exhibited decreased P*_tfoR_* activation and transformation compared to the parent, but this was still significantly greater than Δ*chiS* (**Fig. 3B-D**). Because strains with Chi37 still retained the ability to activate P*_tfoR_* to some degree, it suggests that ChiS-TfoS interactions may not be strictly required for allosteric regulation of TfoS activity. This result is also consistent with our prior observation that TfoS can induce P*_tfoR_* in *E. coli* even in the absence of ChiS [8]. However, these findings also make it clear that the interaction between ChiS and TfoS is important for optimal P*_tfoR_* induction.

**Fig. 3.**
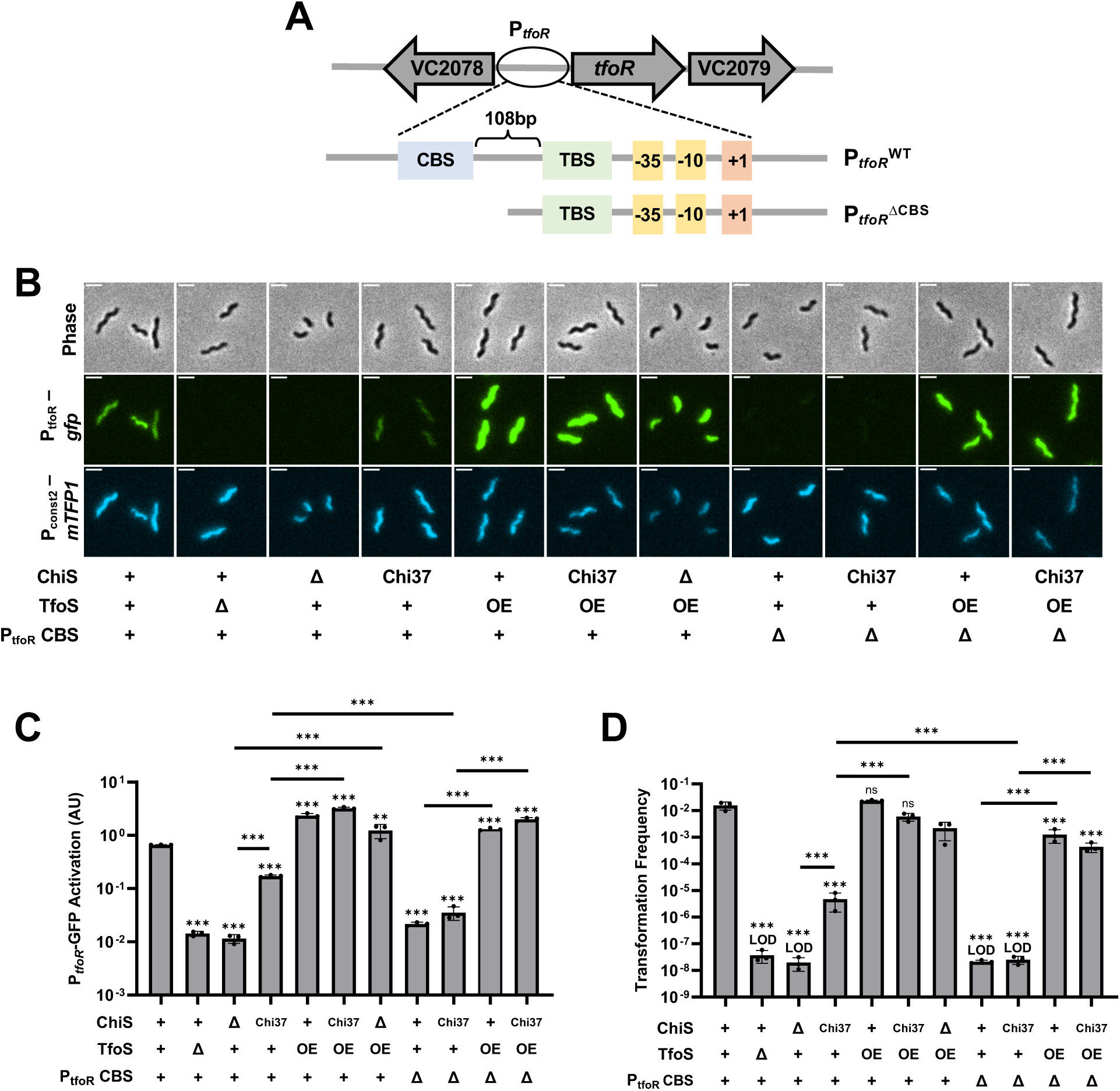
ChiS binding at P_tfoR_ is critical for activation when TfoS is expressed at native levels. Strains harbored P*_tfoR_-gfp,* P*_const2_-mTFP1*, and the indicated mutations. Cells were incubated on chitin and then imaged via epifluorescence microscopy to assess reporter expression. For ChiS genotypes, “+” denotes that cells have a WT copy of ChiS, “Δ” denotes that cells lack ChiS, and “Chi37” indicates that cells express the Chi37 chimera instead of ChiS. For TfoS, “+” denotes TfoS^WT^, “Δ” denotes that TfoS is deleted, and “OE” denotes that TfoS is overexpressed (P*_tac_*-*tfoS +* 1µM IPTG). For P*_tfoR_* CBS, “+” denotes the full-length wildtype P*_tfoR_* promoter while “Δ” denotes P*_tfoR_^ΔCBS^*. (**A**) Schematic of the P*_tfoR_* and P*_tfoR_^ΔCBS^* reporters, highlighting the organization of the ChiS Binding Site (CBS), the TfoS Binding Site (TBS), the –35 and –10 elements, and the transcriptional start site (+1). (**B**) Representative images of cells. Scale bar, 2µm. (**C**) Quantification of P*_tfoR_-gfp* reporter activity. For each replicate (*n* = 3), the geometric mean fluorescence was determined by analyzing 300 individual cells. (**D**) Chitin-dependent transformation assays of the indicated strains. The mutations in P*_tfoR_* were present in both the P*_tfoR_-gfp* reporter construct and the native P*_tfoR_* locus for transcriptional reporter assays, and at the native P*_tfoR_* locus for transformation assays. Results are from three independent biological replicates and shown as the mean ± SD. Statistical comparisons in **C** and **D** are made by one-way ANOVA with Tukey’s multiple comparison test on the log-transformed data (normal distribution confirmed by Shapiro-Wilk test for all samples except “Δ*chiS* TfoS OE” in **D**, which was excluded from the analysis). Statistical identifiers directly above bars represent comparisons to the parent. ns, not significant. *** = *p* < 0.001, ** = *p* < 0.01. LOD, limit of detection.

### Overexpression of TfoS bypasses the need for ChiS to activate P_tfoR_

Because ChiS-TfoS interactions contribute to optimal P*_tfoR_* induction, we hypothesized that this interaction might allow ChiS to recruit TfoS to the membrane-localized P*_tfoR_* DNA locus. Based on this model, we hypothesized that the overexpression of TfoS within the cell should bypass the need for ChiS-TfoS interactions for optimal P*_tfoR_* activation. Under these conditions, we predicted that the likelihood of TfoS being physically close to a membrane-proximal P*_tfoR_* locus should be greatly increased, thereby resulting in restoration of P*_tfoR_* activation in the Chi37 background. To test this hypothesis, we generated an IPTG-inducible TfoS construct (P*_tac_*-*tfoS*) and confirmed that this construct results in overexpression of TfoS via western blot analysis (**Fig. S2C-D**).

Consistent with the proposed model, we found that overexpressing TfoS in the Chi37 background restored P*_tfoR_* activation and natural transformation, suggesting that the ChiS-TfoS interactions were no longer necessary due to the increased abundance of TfoS in the cell (**Fig. 3B-D**). Furthermore, we found that overexpression of TfoS overcomes the need for ChiS completely (*i.e.*, overexpression of TfoS recovers P*_tfoR_* induction and transformation in the Δ*chiS* background) (**Fig. 3B-D**). In contrast, the overexpression of ChiS does not overcome the need for TfoS (**Fig. S5**). These results suggest that the ChiS-dependent recruitment of P*_tfoR_* to the membrane is only required when TfoS is present at native levels; when TfoS is overexpressed, it obviates the need for ChiS by increasing the likelihood that TfoS will come into contact with the P*_tfoR_* DNA locus (which will presumably be membrane-proximal at some frequency due to diffusion of the DNA locus within the cell) independently.

### The ChiS binding site in P_tfoR_ is critical for activation when TfoS is expressed at native levels

Based on the model suggested above, we hypothesize that the ChiS binding site within P*_tfoR_* should be critical for activation when ChiS and TfoS are present at native levels.

Based on the sequence of the <u>C</u>hiS <u>B</u>inding <u>S</u>ites (CBS) in P*_chb_* [4], a putative CBS within P*_tfoR_* was found at the 5’ end of the promoter upstream of the TfoS binding site (**Fig. 3A**). Indeed, truncating the 5’ end of P*_tfoR_* prevents ChiS binding (**Fig. S6**). We refer to this truncated promoter construct as P*_tfoR_^ΔCBS^*. The P*_tfoR_^ΔCBS^* mutation results in minimal P*_tfoR_* activation and inhibits chitin-induced natural transformation when ChiS and TfoS were present at native levels (**Fig. 3B-D**).

Importantly, this strain still contains a functional copy of ChiS, but ChiS can no longer bind at P*_tfoR_*. If the loss of *tfoR* expression in P*_tfoR_^ΔCBS^* is due to a loss of ChiS-dependent recruitment of P*_tfoR_* to the membrane, we would predict that overexpression of TfoS should bypass the impact of the P*_tfoR_^ΔCBS^* mutation (similar to what was observed in Δ*chiS*). Indeed, we found that overexpression of TfoS recovers P*_tfoR_^ΔCBS^* induction and natural transformation (**Fig. 3B-D**). This result also demonstrates that the truncation used to generate P*_tfoR_^ΔCBS^* does not simply diminish the activity of this promoter.

In P*_tfoR_^ΔCBS^*, ChiS binding is eliminated, preventing ChiS-dependent recruitment of the promoter. Removing ChiS DNA recruitment allowed us to further isolate and test the impact of ChiS-TfoS interactions. The ChiS in P*_tfoR_^ΔCBS^* can still bind to TfoS and other DNA loci (*e.g.*, at P*_chb_*), meaning that ChiS binding to TfoS should still be maintained in this background. In contrast, Chi37 exhibits diminished interactions with TfoS in P*_tfoR_^ΔCBS^* and therefore would likely exhibit greatly reduced allosteric activation. We found that there was no increase in P*_tfoR_^ΔCBS^* activation when cells expressed ChiS vs Chi37 (**Fig. 3B-C**). Consistent with this, neither ChiS or Chi37 is capable of supporting chitin-induced natural transformation in the P*_tfoR_^ΔCBS^* background (**Fig. 3D**). These data suggest that ChiS binding at P*_tfoR_* likely plays a more important role in activation than its interaction with TfoS.

### Recruitment of P_tfoR_ to the membrane is sufficient for TfoS-dependent activation

Together, our results suggest that ChiS binding at P*_tfoR_* is more important than ChiS binding to TfoS for activation to occur. These data are consistent with two potential models. First, ChiS binding recruits P*_tfoR_* to the membrane to make it accessible to TfoS. Alternatively, ChiS binding within P*_tfoR_* may alter promoter architecture, which ultimately exposes the TfoS binding site. The distinction between these two models is that for the latter, there would likely be a requirement for native ChiS-CBS interactions within P*_tfoR_* to alter promoter architecture. However, for the former model, we would predict that recruiting P*_tfoR_^ΔCBS^* to the membrane should be sufficient for activation. To test this hypothesis, we took advantage of the Tet repressor (TetR), which naturally binds its operator sequence, *tetO*, at high affinity. Specifically, we replaced the CBS within P*_tfoR_* with *tetO* (P*_tfoR_^ΔCBS::tetO^*) and swapped the native DNA-binding domain (DBD) of ChiS [23] for the full TetR protein (**Fig. 4A**). When we expressed ChiS^ΔDBD^-TetR in cells containing P*_tfoR_^ΔCBS::tetO^*, we found that induction and natural transformation were partially recovered when compared to cells lacking the ChiS^ΔDBD^-TetR construct (**Fig. 4B-D**). To test this further, we incubated strains with anhydrotetracycline (ATc) to inhibit the DNA-binding activity of TetR. Under these conditions, both P*_tfoR_* induction and natural transformation were ablated. This result is consistent with our hypothesis and suggests that recruitment of P*_tfoR_^ΔCBS::tetO^* to the membrane via ChiS^ΔDBD^-TetR binding is necessary for activation to occur (**Fig. 4B-D**). This result also argues against ChiS-dependent remodeling of *tfoR* promoter architecture, as native ChiS-CBS interactions have been entirely replaced by those of TetR and *tetO*. Importantly, overexpression of TfoS was able to rescue P*_tfoR_^ΔCBS::tetO^* induction even in the absence of ChiS, indicating that this promoter is fully functional (**Fig. 4B-D**).

**Fig. 4.**
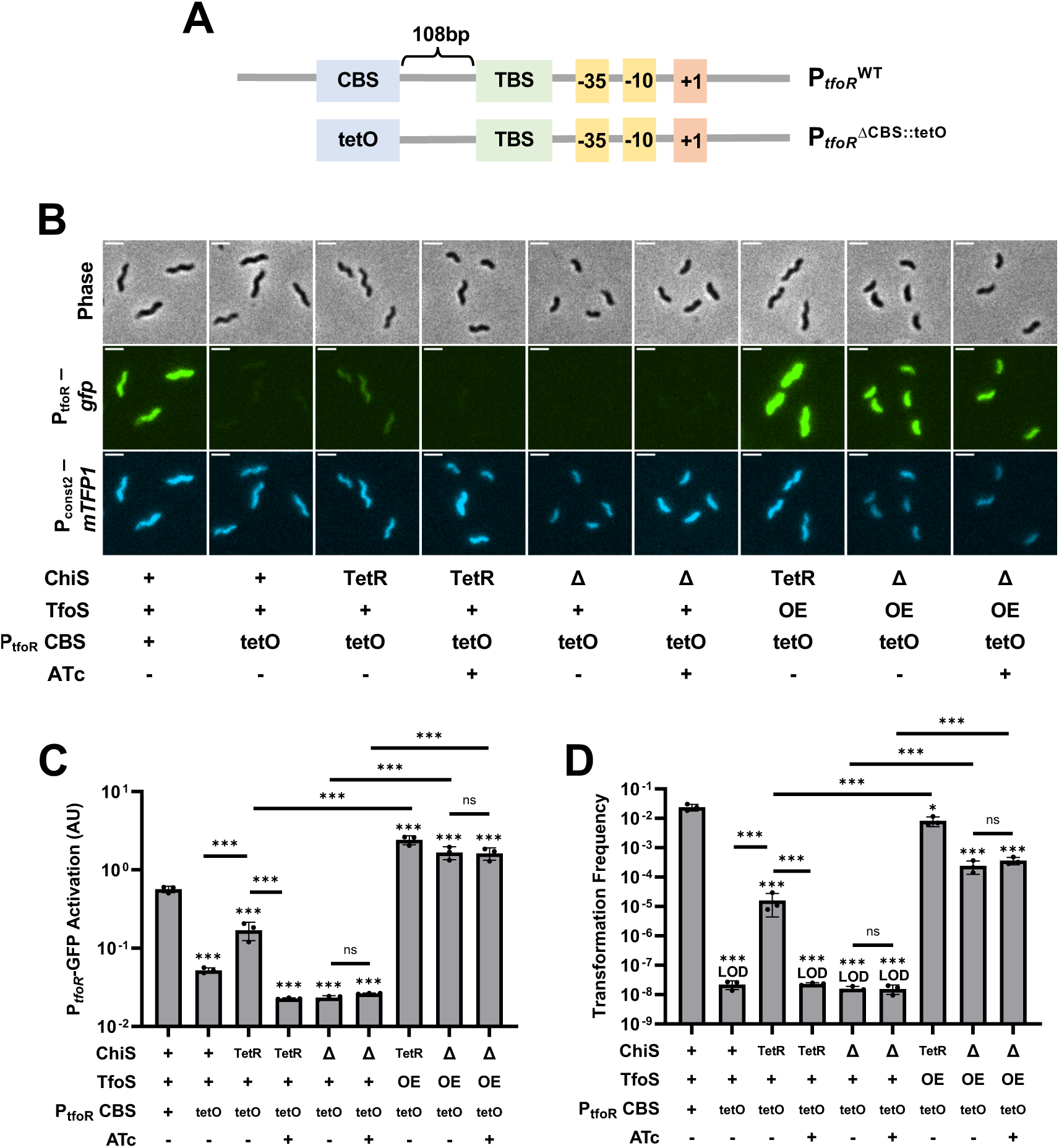
Recruitment of P_tfoR_ to the membrane is sufficient for TfoS-dependent activation. Strains harbored P*_tfoR_-gfp,* P*_const2_-mTFP1*, and the indicated mutations. Cells were incubated on chitin and then imaged via epifluorescence microscopy to assess reporter expression. For ChiS genotypes, “+” denotes that cells have a WT copy of ChiS, “Δ” denotes that cells lack ChiS, and “TetR” indicates that cells express a ChiS^ΔDBD^-TetR fusion. For TfoS, “+” denotes TfoS^WT^, “Δ” denotes that TfoS is deleted, and “OE” denotes that TfoS is overexpressed (P*_tac_*-*tfoS +* 1µM IPTG). For P*_tfoR_* CBS, “+” denotes the full-length wildtype P*_tfoR_* promoter while “tetO” denotes P*_tfoR_^ΔCBS::tetO^*. Where indicated, 50 ng/mL ATc was added to the reaction. (**A**) Schematic of the P*_tfoR_* and *P_tfoR_^ΔCBS::tetO^* reporters, highlighting the organization of the ChiS Binding Site (CBS), the TfoS Binding Site (TBS), the Tet operator (tetO) site, the –35 and –10 elements, and the transcriptional start site (+1). (**B**) Representative images of cells. Scale bar, 2µm. (**C**) Quantification of P*_tfoR_-gfp* reporter activity. For each replicate (*n* = 3), the geometric mean fluorescence was determined by analyzing 300 individual cells. (**D**) Chitin-dependent transformation assays of the indicated strains. The mutations in P*_tfoR_* were present in the P*_tfoR_-gfp* reporter construct for transcriptional reporter assays, but at the native locus for transformation assays. Results are from three independent biological replicates and shown as the mean ± SD. Statistical comparisons in **C** and **D** are made by one-way ANOVA with Tukey’s multiple comparison test on the log-transformed data (normal distribution confirmed by Shapiro-Wilk test). Statistical identifiers directly above bars represent comparisons to the parent. ns, not significant. *** = *p* < 0.001, ** = *p* < 0.01. LOD, limit of detection.

If ChiS binding to P*_tfoR_* primarily serves to recruit the promoter to the membrane, then we would hypothesize that the helical phasing of the CBS to the TfoS binding site (TBS) within P*_tfoR_* is not critical for activation. To test this, we inserted either 5 bp (loss of phasing) or 10 bp (retains phasing) between the CBS and TBS in P*_tfoR_* (**Fig. S7**).

Inserting 5 bp, which disrupts phasing between the CBS and TBS, resulted in almost parent levels of P*_tfoR_* activation. Similarly, insertion of 10 bp, which maintains phasing, resulted in full activation of P*_tfoR_* (**Fig. S7**). Together, these results suggest that neither the phasing between the CBS and TBS nor the exact position of the CBS in P*_tfoR_* is critical for activation. These results are consistent with a model in which ChiS binding to P*_tfoR_* primarily serves to relocalize this promoter to the membrane.

## DISCUSSION

In this study, we uncover how ChiS and TfoS coordinate to promote activation of natural transformation in *V. cholerae*. Specifically, our data are most consistent with a model wherein ChiS performs two functions to facilitate P*_tfoR_* activation: (1) it recruits the P*_tfoR_* DNA locus to the membrane and (2) it recruits TfoS to the membrane-proximal promoter through a direct interaction (**Fig. 5**).

**Fig. 5.**
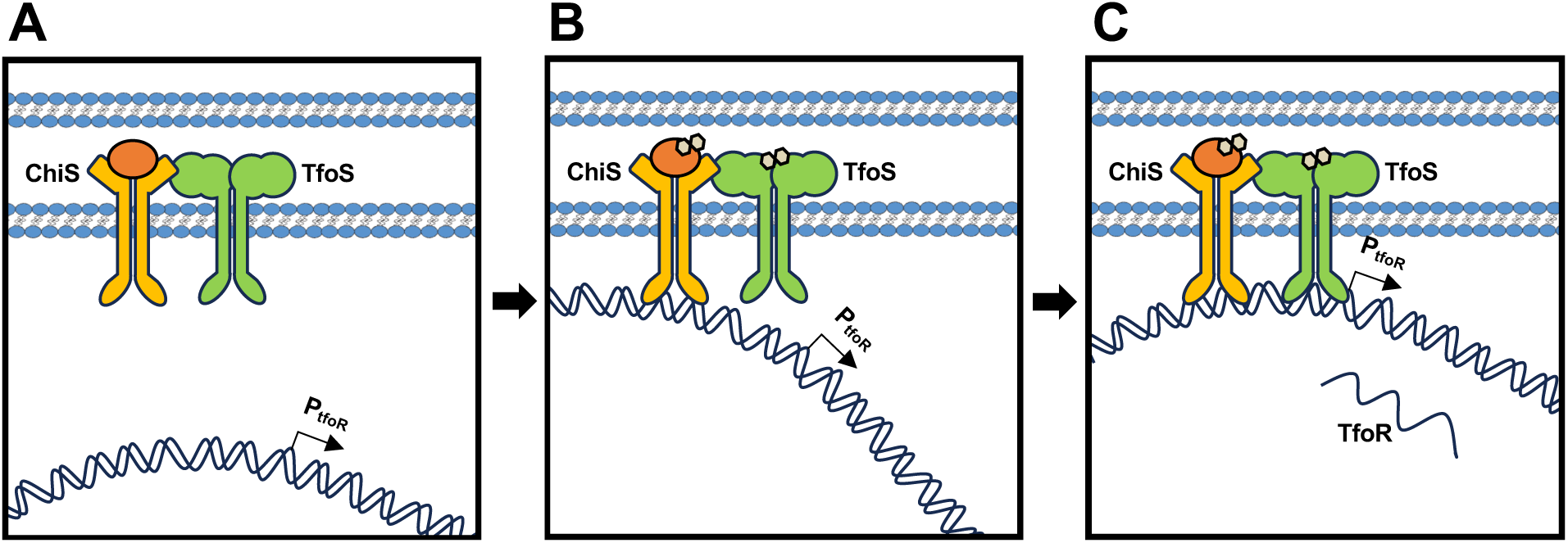
Proposed model for chitin-induced activation of P_tfoR_. (**A**) In the absence of chitin oligosaccharides (tan hexagons), ChiS and TfoS are inactive. In the presence of chitin, (**B**) ChiS and TfoS are activated and ChiS binds upstream of the *tfoR* promoter. ChiS binds to TfoS, thereby recruiting it to the membrane-localized P*_tfoR_*. (**C**) These combined activities of ChiS allow TfoS to subsequently bind P*_tfoR_* and activate transcription of the *tfoR* small RNA that is critical for chitin-induced natural transformation.

The downstream effect of *tfoR* expression is the activation of the genes required for DNA uptake and integration by natural transformation. It is hypothesized that natural transformation is induced by chitin oligosaccharides because *Vibrio* species accumulate and form biofilms on chitinous surfaces in the aquatic environment. Thus, chitin is a niche where diverse *Vibrios* are present, and represents an ideal site for genetic exchange. In support of this model, prior work has uncovered that on chitin, *V. cholerae* actively lyses neighboring bacteria to release their DNA as a substrate for natural transformation [24, 25].

Our data suggests that simply overexpressing TfoS is sufficient to bypass the requirement for ChiS for P*_tfoR_* activation. ChiS and TfoS are likely independently activated by chitin oligosaccharides [4, 8]. Why then has ChiS-dependent regulation of P*_tfoR_* been maintained in *V. cholerae*? One possibility is that this dual requirement ensures that natural transformation is not induced until cells are experiencing conditions where chitin oligosaccharides are abundant. This would prevent the inappropriate expression of the large repertoire of genes required for natural transformation, which might incur a significant fitness cost. This regulation is reminiscent of the “coincidence detector” described for quorum sensing in *Vibrio* spp., which requires the simultaneous detection of multiple autoinducers before activation can occur [26].

However, not all genes within the chitin regulon require the dual activity of ChiS and TfoS. For example, the *chb* operon required for chitin uptake and catabolism is largely dependent on ChiS alone as shown in this and prior work [4, 9]. Because nutrient acquisition is likely a prerequisite for natural transformation, it is tempting to speculate that the “relaxed” requirement for chitin-dependent induction of the *chb* operon is to ensure that chitin uptake and catabolism is prioritized over genetic exchange by natural transformation.

Here, we show that ChiS and TfoS are two TTRs that must cooperatively act to promote P*_tfoR_* activation. This cooperation between TTRs may be a common feature for this class of transcriptional regulators. Indeed, *V. cholerae* contains another pair of TTRs, ToxRS and TcpPH, that coordinate to activate the *toxT* promoter [5–7, 27, 28]. There are a number of parallels between ToxRS/TcpPH dependent activation of P*_toxT_* and ChiS/TfoS dependent activation of P*_tfoR_*. Namely, TcpPH is sufficient to activate P*_toxT_* when overexpressed, but requires ToxRS for P*_toxT_* induction when expressed at native levels [27]. Furthermore, the membrane-embedded localization of ToxRS is required for this stimulatory activity [7, 28]. The ToxRS binding site in P*_toxT_* is also located upstream of the TcpPH binding site [7, 27]. These data are consistent with ToxRS facilitating recruitment of P*_toxT_* to the membrane for downstream binding and activation by TcpPH. Recent results, however, suggest that ToxRS interactions with P*_toxT_* may alter the structure of the promoter to promote TcpPH-dependent activation [5, 6]. TcpPH and ToxRS are members of a rapidly evolving family of signal transduction proteins in enteric bacteria [29]. Thus, other members of this family may also coordinate to regulate their downstream targets. One recent example is VrtAC, which likely coordinates with VtrB to activate the expression and transertion of the type III secretion system in *V. parahaemolyticus* [3]. Not all TTRs require cooperativity for their activity, however. This is true for ChiS at the *chb* promoter [4, 9], ToxR at the *ompU* promoter [27], and *E. coli* CadC at the *cadBA* operon [2].

The localization and diffusion of TTRs is clearly constrained compared to canonical cytoplasmic TRs. However, whether this constraint impacts how TTRs bind to their target sites is not clear. Our results suggest that natively expressed TfoS cannot activate P*_tfoR_* unless ChiS is also present and able to bind P*_tfoR_*. However, this requirement for ChiS can be overcome by simply overexpressing TfoS. This finding suggests that when natively expressed, TfoS cannot bind to P*_tfoR_* in a manner that is sufficient to promote activation. Thus, the accessibility of the DNA binding targets for membrane-constrained TTRs may represent an additional layer of regulation that is a distinct property for this class of regulators when compared to canonical cytoplasmic transcriptional regulators. The DNA targets that TTRs bind must localize proximally to the membrane at some frequency. However, it is unclear if all regions of the genome are equally likely to diffuse and localize near the membrane in this manner. One possibility is that TTRs have evolved to bind to genomic loci that are more likely to come into contact with the membrane. Future work will focus on systematically assessing whether the constraint of membrane localization limits the genomic loci that TTRs can effectively bind and regulate.

Our data are consistent with a model in which membrane recruitment of P*_tfoR_* is necessary for TfoS-dependent activation. This could be because TfoS, when expressed at native levels, lacks the ability to bind and recruit P*_tfoR_* to the membrane by itself. However, the P*_tfoR_* locus is not inherently difficult for a TTR to bind because we show that ChiS naturally binds to this locus from the membrane. Therefore, the P*_tfoR_* DNA locus must come into contact with the membrane at some frequency. If so, one interpretation of our data is that TfoS is less capable of stably binding P*_tfoR_* at the membrane compared to ChiS. Indeed, kinetic studies of the TTR CadC suggest that the mobility of the CadC protein within the membrane, but not the mobility of its DNA target, is rate-limiting for CadC-DNA interactions [30]. This finding indicates that some regions of the genome may contact the membrane with sufficient frequency that their diffusion / localization does not limit the TTR-DNA target search.

DNA-binding transcriptional regulators spend a large portion of their time transiently binding to DNA nonspecifically during their target search [31]. This transient binding can keep these proteins in close proximity to the nucleoid and help reduce the dimensionality of the 3D target search for a freely diffusing cytoplasmic DNA-binding protein. Therefore, a distinct possibility is that the membrane localization of TfoS does not make it more difficult for this regulator to bind to DNA. Instead, it could be that TfoS is capable of binding DNA, but does so with a relaxed specificity, which reduces the likelihood that it will identify and bind its sequence-specific target sites within the genome compared to other TTRs like ChiS. If so, ChiS binding at P*_tfoR_* may simply “trap” this DNA locus at the membrane and help reduce the dimensionality of the target search for TfoS from a three-dimensional search into a two-dimensional search (due to the constrained diffusion of TfoS within a planar membrane). It is tempting to speculate that other molecular interactions, whether protein-protein or DNA-protein, also take advantage of the membrane as a planar space to increase the specificity and/or efficiency of the target search.

## METHODS

### Bacterial strains and culture conditions

All mutant strains were derived from the *Vibrio cholerae* El Tor strain E7946 [32]. Unless otherwise indicated, *V. cholerae* strains were grown in LB medium or on LB agar. When necessary, LB was supplemented with chloramphenicol (1 µg/mL), kanamycin (50 µg/mL), trimethoprim (10 µg/mL), spectinomycin (200 µg/mL), carbenicillin (100 µg/mL), erythromycin (10 µg/mL), zeocin (100 µg/mL), or sulfamethoxazole (100 µg/mL). A list of all strains used in this study can be found in **Table S1**.

### Strain construction

Mutant constructs were made using splicing-by-overlap extension (SOE) PCR, as previously described [33]. Chromosomally integrated ectopic expression constructs were used throughout this study as previously described [34]. A list of all primers used to make mutant constructs can be found in **Table S2**. Mutant strains were generated by introducing constructs into strains by natural transformation; either chitin-induced natural transformation or chitin-independent natural transformation via the use of an IPTG-inducible *tfoX* plasmid (pMMB-*tfoX*) [35]. The pMMB-*tfoX* plasmid was cured before strains were used in any experiments. Mutants were confirmed by PCR and/or sequencing.

### Chitin-induced natural transformation assays

Strains were grown overnight in LB broth at 30°C rolling. Overnight cultures were then subcultured in LB broth, with 1 µM IPTG when applicable, and grown to mid-log phase (OD_600_ ∼0.5-1.0). Cells were washed and resuspended in 0.5X Instant Ocean (IO; 7g/L, Aquarium Systems Inc.). Then, 1 mL chitin reactions were made by mixing 10^8^ cells (suspending in 100 µL IO), 150 µL of chitin slurry (8g/150mL, Alfa Aesar), 750 µL IO, and 1 µM IPTG when applicable. Chitin reactions were incubated at 30°C static for ∼18 hours. Next, 550 µL of supernatant was removed, and 500 ng of transforming DNA (tDNA) was mixed with the cells by inversion. The tDNA used replaced the frame-shifted transposase gene VC1807 (*i.e.*, a “neutral” gene locus) with either an Erm^R^ or Cm^R^ marker. A no tDNA control was included for each strain tested. Cells were incubated with DNA for 5 hours at 30°C static. Then, 500 µL LB broth was added to each reaction, and reactions were outgrown by shaking at 37°C for 2 hours. After outgrowth, reactions were plated for quantitative culture for total viable count (plated on plain LB agar) and transformants (plated on LB agar with antibiotic). Transformation frequency was calculated by dividing the CFU/mL of transformants by the CFU/mL of total viable counts.

### Microscopy

Cells were imaged on an inverted Nikon Ti-2 microscope with a Plan Apo 60x objective lens, the appropriate filter cubes (FITC, YFP, mCherry, and/or CFP), a Hamamatsu ORCAFlash 4.0 camera, and Nikon NIS Elements imaging software. Image analysis was conducted using the MicrobeJ plugin [36] in Fiji [37]. Cell body segmentation was performed using the “medial axis” mode on the phase image. Segmentation parameters were adjusted for each image, but generally fell within the following ranges: minimum area of 0.45 µm^2^, length between 0.6-2.7 µm, and width between 0.3-1 µm. Fluorescence was measured in the interior of segmented cells, which included the cell periphery and cytoplasm.

### PopZ-linked apical recruitment (POLAR) assays

Strains were grown overnight in LB broth at 30°C rolling and were supplemented with 10 µM IPTG to induce *P_tac_-tfoS-mCherry, P_tac_-chiS-GFP-H3H4, and/or P_tac_-chi37-GFP-H3H4* constructs. For POLAR experiments, LB broth was also supplemented with 0.05% arabinose to induce *P_bad_-PopZ* expression. For efficient polar relocalization and to avoid issues with cell curvature, the curvature determinant *crvA* [38] was deleted in all colocalization strains. Overnight cultures were washed and resuspended in 0.5X IO. Cells were then placed on a glass coverslip beneath a 0.4% IO gelzan pad and imaged using FITC and mCherry filter cubes. Heat maps were generated by analyzing the fluorescence localization of 300 cells per replicate. To quantify polar localization, the distance of fluorescence to the nearest cell pole was measured in each channel. Fluorescence within 0.2 µm of the cell pole was conservatively denoted as polarly localized.

### Bacterial adenylate cyclase two-hybrid assays

ChiS and TfoS genes were amplified using PCR and cloned into the BACTH vectors pKT25, pUT18C, pKNT25, and pUT18 to generate N– and C-terminal fusions to the T18 and T25 fragments of adenylate cyclase. Miniprepped plasmids were then cotransformed into *E. coli* BTH101. Transformations were plated onto LB agar plates with kanamycin (50 µg/mL) and carbenicillin (100 µg/mL) to select for transformants that received both plasmids. The resulting strains were then grown up overnight in LB broth at 30°C, and 3 µL of the overnight culture was spotted onto an LB agar plate supplemented with 500 μM IPTG, kanamycin (50 μg/mL), carbenicillin (100 μg/mL), and 5-bromo-4-chloro-3-indolyl-β-d-galactopyranoside (X-gal) (40 μg/mL). Plates were incubated at 30°C for 48 hours before imaging on an HP Scanjet G4010 flatbed scanner.

### Electrophoretic mobility shift assays (EMSAs)

Binding reactions contained 10 mM Tris·HCl pH 7.5, 1 mM EDTA, 10 mM KCl, 1 mM DTT, 50 µg/mL BSA, 0.1 mg/mL salmon sperm DNA, 5% glycerol, 3 nM DNA probe, and purified MBP-ChiS cytoplasmic domain [4] at 25 nM, 50 nM, 100 nM, 200 nM, 400 nM, and 800 nM concentrations (diluted in 10 mM Tris pH 7.5, 10 mM KCl, 1 mM DTT, and 5% glycerol). Binding reactions were incubated in the dark at room temperature for 20 minutes before being loaded into a 0.5X Tris Borate EDTA (TBE) polyacrylamide gel. Gels were run in 0.5X TBE buffer at 4°C, then imaged for Cy5 fluorescence on a ProteinSimple FluorChem R system. Cy5-labeled DNA probes were made by Taq PCR with 4.34 µM Cy5-dCTP. Unlabeled probes were made by Phusion PCR.

### Fluorescence reporter assays

Strains were grown overnight in LB broth, supplemented with 1 µM IPTG and/or 50 ng/mL ATc when applicable, at 30°C rolling. Then, overnight cultures were washed and resuspended in IO. 1 mL chitin reactions were made by mixing 10^8^ cells (suspended in 100 µL IO), 150 µL of chitin slurry (8g/150mL), 750 µL IO, and 1 µM IPTG and/or 50 ng/mL ATc when applicable. Chitin reactions were incubated at 30°C shaking for 24 hours. Next, reactions were vortexed to dislodge the cells from the chitin. The supernatants (containing the resuspended cells) were then centrifuged, resuspended in IO, and placed on a glass coverslip beneath a 0.2% IO gelzan pad. Cells were then imaged using mCherry, YFP, and CFP filter cubes. Image analysis was performed to quantify the average fluorescence per cell in each channel. The background fluorescence was determined in each channel by imaging and analyzing a non-fluorescent strain, which was then subtracted from all samples. Reporter fluorescence (GFP or mCherry) was then divided by the fluorescence of the CFP channel to normalize against the constitutively expressed mTFP1 construct. The geometric mean was then calculated from 300 individual cells for each replicate.

### Western blotting

Strains were grown overnight in LB broth, supplemented with 1 µM IPTG when applicable, at 30°C rolling. Overnight cultures were then washed and resuspended to OD_600_=100 in IO. Cells were then mixed 1:1 with 2X SDS-PAGE sample buffer (200 mM Tris pH 6.8, 25% glycerol, 4.1% SDS, 0.02% Bromophenol Blue, 5% β-mercaptoethanol) and boiled for 10 minutes. To blot for TfoS-FLAG, 4 µL of each sample was separated on a 15% SDS-PAGE gel. To blot for RpoA, 2 µL of each sample was separated on a 15% SDS-PAGE gel. Following SDS-PAGE, the proteins were transferred to a polyvinylidene difluoride (PVDF) membrane by electrophoresis, and then membranes were blocked for 1 hour in 2% milk. Next, membranes were incubated rocking at room temperature overnight with primary antibody: rabbit polyclonal anti-FLAG (Sigma) or mouse monoclonal anti-RpoA (Biolegend). Membranes were then washed and incubated with either anti-mouse or anti-rabbit horseradish peroxidase-conjugated secondary antibody for three hours rocking at room temperature. Then, blots were washed, developed with Pierce ECL Western blotting substrate, and imaged with a ProteinSimple FluorChem R system.

### Chromatin immunoprecipitation assays

Chromatin Immunoprecipitation (ChIP) assays were performed exactly as previously described [39]. *V. cholerae* strains were initiated from freezer stocks and incubated at 30°C in LB broth rolling for ∼16 hours. The following day, cultures were subcultured into fresh LB to an OD_600_ of 0.08 and incubated rolling for 6 hours at 30°C. Cells were incubated with 1% paraformaldehyde at room temperature for 20 minutes shaking to crosslink DNA and proteins. A 1.2 molar excess of Tris (10 minutes, room temperature, shaking) was used to quench residual paraformaldehyde. Crosslinked cells were washed twice with TBS (25 mM Tris HCl, pH 7.5 and 125 mM NaCl) and then frozen at – 80°C. Cells were then resuspended in lysis buffer (1x FastBreak cell lysis reagent (Promega), 50 μg/mL lysozyme, 1% Triton X-100, 1 mM PMSF, and 1x protease inhibitor cocktail; 100x inhibitor cocktail contained the following: 0.07 mg/mL phosphoramidon (Santa Cruz), 0.006 mg/mL bestatin (MPbiomedicals/Fisher Scientific), 1.67 mg/mL AEBSF (Gold Bio), 0.07 mg/mL pepstatin A (DOT Scientific), 0.07 mg/mL E64 (Gold Bio) suspended in DMSO) to an OD_600_ of 50.0. These resuspensions were incubated rocking at room temperature for 20 minutes to permit cell lysis. Lysates were then sonicated to shear DNA into 200-300bp fragments (sonication was performed with a Fisherbrand Model 705 Sonic Dismembrator 6 times for 10 seconds at 1% amplitude, resting on ice for at least 30 seconds between each sonication). The sonicated lysates were then clarified by centrifugation and diluted 1:5 in Immunoprecipitation (IP) Buffer (50 mM HEPES NaOH pH 7.5, 150 mM NaCl, 1 mM EDTA, and 1% Triton X-100). An “input DNA” sample was collected from this diluted lysate. 1 mL of this diluted lysate was then added to Pierce anti-DYKDDDDK magnetic agarose equilibrated in IP buffer and then incubated at room temperature with end-over-end mixing. After two hours of incubation, a magnetic separation rack was used to facilitate removal of the supernatant. The agarose was washed twice with IP Buffer for 1 minute, once with Wash Buffer 1 (50 mM HEPES NaOH pH 7.5, 1 mM EDTA, 1% Triton X-100, 500 mM NaCl, and 0.1% SDS) for 5 minutes, once with Wash Buffer 2 (10 mM Tris HCl pH 8.0, 250 mM LiCl, 1 mM EDTA, 0.5% NP-40, and 1% Triton X-100) for 5 minutes, and once with Buffer TE (10 mM tris pH 8.0 and 1 mM EDTA) for 1 minute. SDS elution buffer (50 mM Tris HCl pH 7.5, 10 mM EDTA, and 1% SDS) was used to elute protein and DNA off the magnetic agarose (30 minutes at 65°C). Supernatant was collected and digested for 2 hours with 20 μg Proteinase K in SDS Elution Buffer at 42°C. 6 hours of incubation at 65°C was then used to reverse crosslinking. Both the input and the output samples were then purified to remove non-DNA components. The abundance of P*_tfoR_* in the input DNA (diluted 1:100) and output DNA was then quantified using quantitative PCR with iTaq Universal SYBR Green Supermix (Bio-Rad). A fold enrichment value was then generated by measuring the abundance of the *tfoR* promoter in the output (following immunoprecipitation) relative to the input (prior to immunoprecipitation). A locus that is not bound by ChiS or TfoS (*rpoB*) was used to normalize for total amount of DNA in the sample.

### ChIP-seq assays

ChIP input and output DNA were prepared exactly as described in “Chromatin immunoprecipitation assays” above. Sequencing libraries were prepared using the NEBNext Ultra II DNA Library Prep Kit (New England Biolabs) for Illumina according to manufacturer’s instructions. Libraries were then analyzed on a 4200 TapeStation (Agilent), pooled, and then loaded on a NextSeq 1000/2000 P2 Reagents (100 Cycles) v3 flow cell configured to generate 2 x 61 nt paired-end reads. Demultiplexing was performed with bcl2fastq, version 2.20.0.

Reads were then analyzed using the open-source web-based Galaxy server [40]. Demultiplexed paired-end fastq files were aligned to the *V. cholerae* genome (*Vibrio cholerae* O1 biovar El Tor str. N16961, NCBI RefSeq assembly: GCF_000006745.1) using bowtie2 with default parameters for paired-end data. Output BAM files were converted to bigWig files for peak visualization using bamCoverage. Bin size was set to 50 bp and coverage was normalized to 1x. Effective genome size was user-specified at 4,033,464. The tool bigwigCompare was then used to visualize the enrichment of output over input bigWig files. The ChIP output file was set as the “treatment file” and the ChIP input was set as the “bigwig file” and the option “compute the ratio signals” was used to generate plots for of the ChIP ratio. To account for regions of zero coverage, a pseudocount of 0.01 was added to each binned region. The output bigWig file was then visualized on Integrated Genome Viewer 2.8.4. Raw and processed data were uploaded to NCBI GEO (Accession: GSE284091).

### Statistical Comparisons

All statistical comparisons were made in Graphpad Prism and the statistical tests used are indicated in figure legends. For summary statistics and a complete list of all statistical comparisons, see **S1 Dataset**.

## ACKNOWLEDGEMENTS

We would like to thank the Center for Genomics and Bioinformatics at Indiana University for assistance with ChIP-seq library prep and sequencing. We would also like to thank Gabe Zentner for his assistance and advice on ChIP-seq analysis. This work was supported by grant R35GM128674 from the National Institutes of Health to ABD.

---

**Fig. S1.**
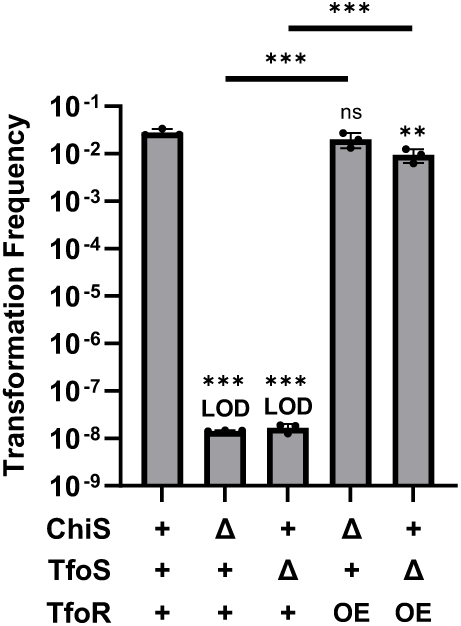
Overexpression of tfoR recovers transformation in ΔchiS and ΔtfoS backgrounds. Chitin-dependent transformation assays of the indicated strains. For ChiS genotypes, “+” denotes that cells have a WT copy of ChiS and “Δ” denotes that cells lack ChiS. For TfoS genotypes, “+” denotes TfoS^WT^ and “Δ” denotes that TfoS is deleted. For *tfoR*, “+” denotes WT *tfoR* and “OE” denotes that *tfoR* is overexpressed (P*_tac_-tfoR* + 1µM IPTG). Results are from three independent biological replicates and shown as mean ± SD. Statistical comparisons are made by one-way ANOVA with Tukey’s multiple comparison test on the log-transformed data (normal distribution confirmed by Shapiro-Wilk test). Statistical identifiers directly above bars represent comparisons to the parent. ns, not significant. *** = *p* < 0.001, ** = *p* < 0.01. LOD, limit of detection.

**Fig. S2.**
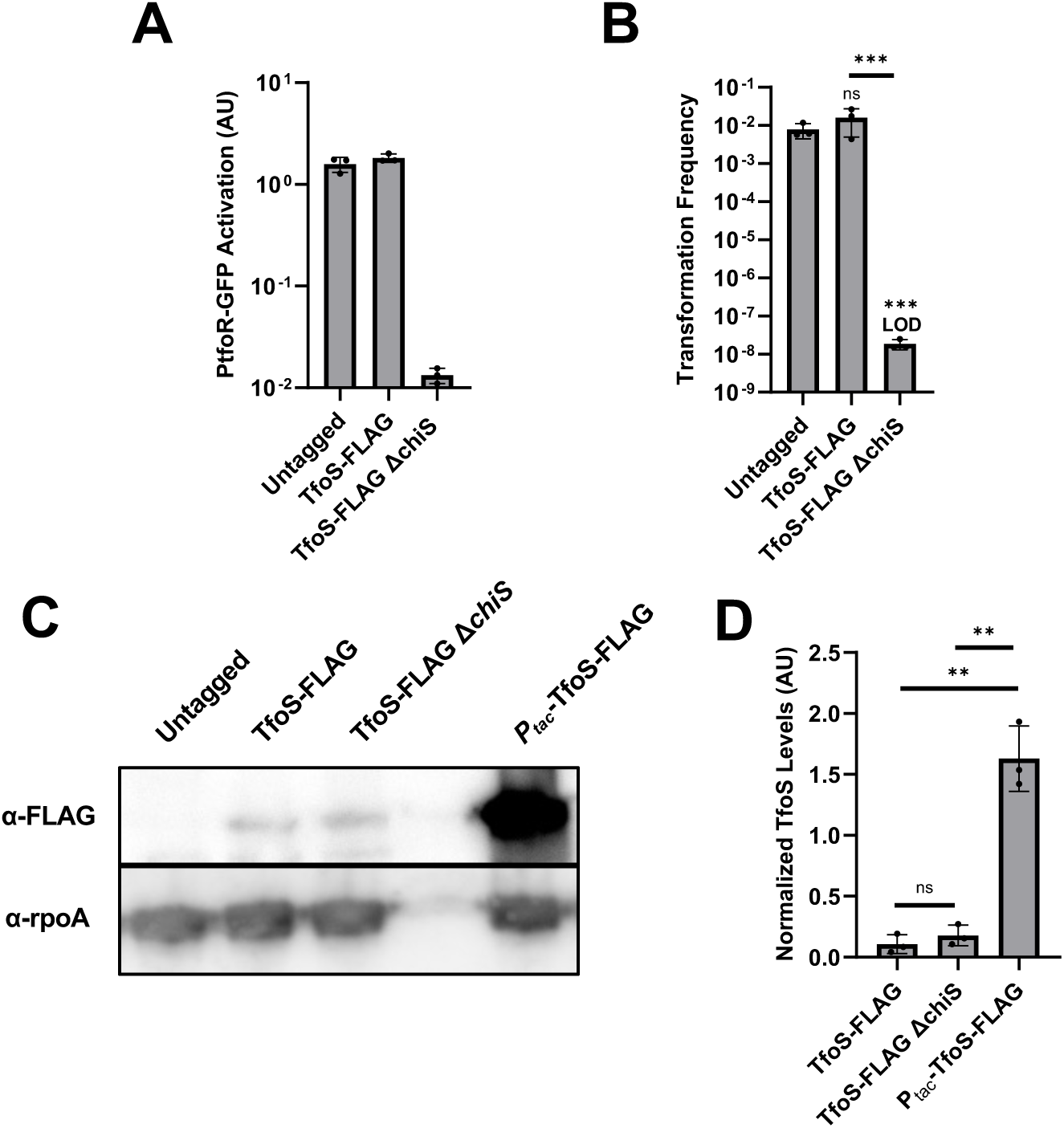
ChiS does not alter TfoS expression. (**A**) Transcriptional reporter assays to assess chitin-dependent gene expression in the indicated strains. All strains harbored P*_tfoR_-gfp* and P*_const2_-mTFP1* constructs. Cells were incubated on chitin and then imaged via epifluorescence microscopy to assess reporter expression. For each replicate (*n* = 3), the geometric mean fluorescence was determined by analyzing 300 individual cells. (**B**) Chitin-dependent transformation assays of indicated strains. (**C**) Western blot analysis of the indicated strains. The blot in **C** is representative of three biological replicates. (**D**) Quantification of TfoS levels from western blots. For each replicate (*n* = 3), the normalized TfoS level was determined by dividing the intensity of the TfoS band by the intensity of the RpoA band. Results in **A**, **B**, and **D** are from three independent biological replicates and shown as the mean ± SD. Statistical comparisons in **B** and **D** were made by one-way ANOVA with Tukey’s multiple comparison test (normal distribution confirmed by Shapiro-Wilk test). Statistical identifiers directly above bars represent comparisons to the parent. ns, not significant. *** = *p* < 0.001, ** = *p* < 0.01. LOD, limit of detection.

**Fig. S3.**
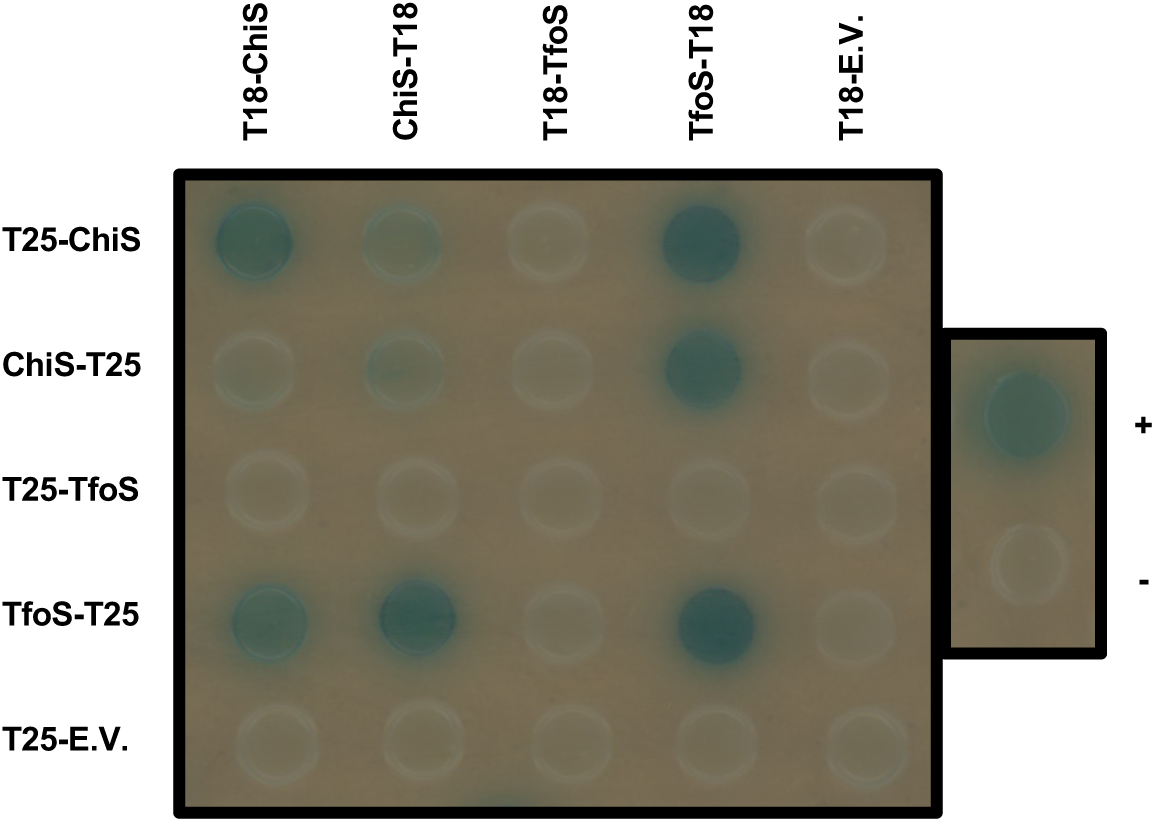
ChiS interacts with TfoS. BACTH assess interactions between ChiS and TfoS. For BACTH assays, ChiS and TfoS either had an N-terminal fusion (indicated as TXX-protein) or a C-terminal fusion (indicated as protein-TXX) to the T25 or T18 fragment of adenylate cyclase.“+” and “-” indicate positive (T18-zip + T25-zip) and negative (T18 E.V. + T25 E.V.) controls for the assay. “E.V.” denotes an empty vector.

**Fig. S4.**
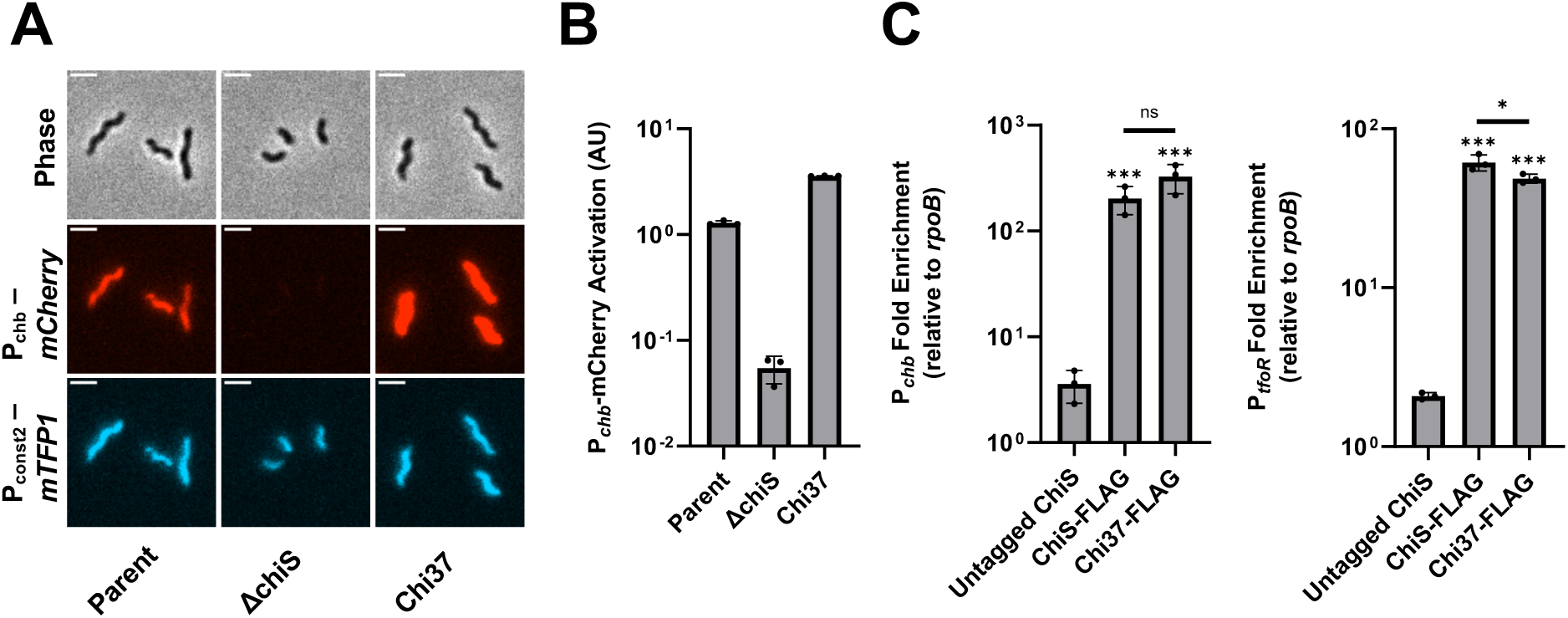
Chi37 is a functional chimera of ChiS. (**A**-**B**) Transcriptional reporter assays to assess chitin-dependent gene expression in the indicated strains. All strains harbored P*_chb_-mCherry* and P*_const2_-mTFP1* constructs. Cells were incubated on chitin and then imaged via epifluorescence microscopy to assess reporter expression. Representative images are shown in **A** and the quantification of the results are shown in **B**. For each replicate (*n* = 3), the geometric mean fluorescence was determined by analyzing 300 individual cells. (**C**) ChIP-qPCR assays were performed with the indicated strains to assess ChiS and Chi37 binding at P*_chb_* and P*_tfoR_ in vivo*. Results are from three independent biological replicates and shown as the mean ± SD. Statistical comparisons were made by one-way ANOVA with Tukey’s multiple comparison test (normal distribution confirmed by Shapiro-Wilk test). Statistical identifiers directly above bars represent comparisons to the parent. ns, not significant. *** = *p* < 0.001, * = *p* < 0.05.

**Fig. S5.**
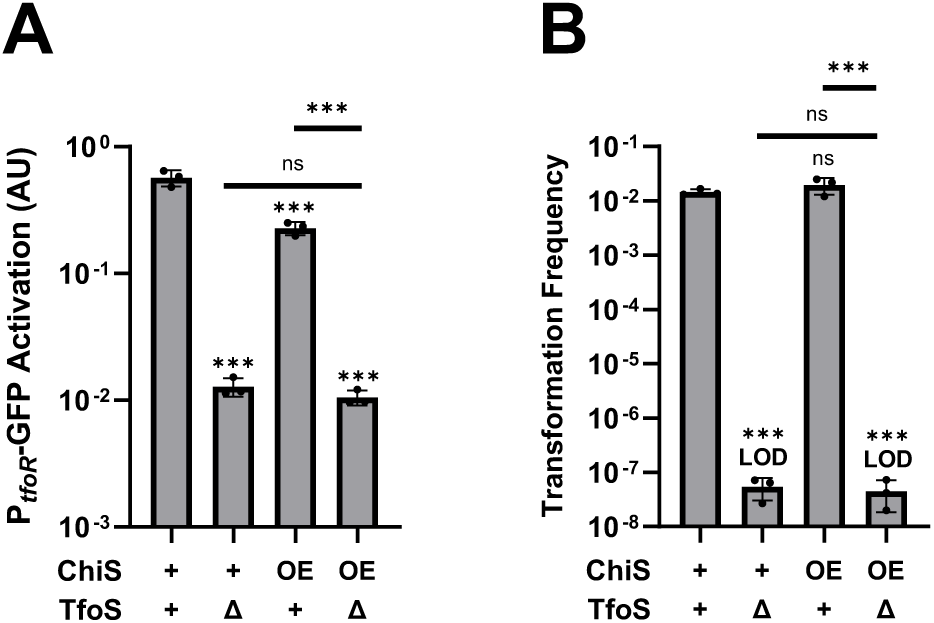
Overexpression of ChiS does not restore P_tfoR_ activation in the absence of TfoS. (**A**) Transcriptional reporter assays to assess chitin-dependent gene expression in the indicated strains. For ChiS genotypes, “+” denotes cells that have a WT copy of ChiS, and “OE” denotes that ChiS is overexpressed (P*_tac_-chiS* + 1µM IPTG). For TfoS genotypes, “+” indicates that cells have a WT copy of TfoS, while “Δ” denotes that cells lack TfoS. All strains harbored P*_tfoR_-gfp* and P*_const2_-mTFP1* constructs. For each replicate (*n* = 3), the geometric mean was determined by analyzing 300 individual cells. (**B**) Chitin-dependent transformation assays of the indicated strains. Results are from three independent biological replicates and shown as mean ± SD. Statistical comparisons were made by one-way ANOVA with Tukey’s multiple comparison test on the log-transformed data (normal distribution confirmed by Shapiro-Wilk test). Statistical identifiers directly above bars represent comparisons made to the parent. *** = *p* < 0.001, ns = not significant. LOD, limit of detection.

**Fig. S6.**
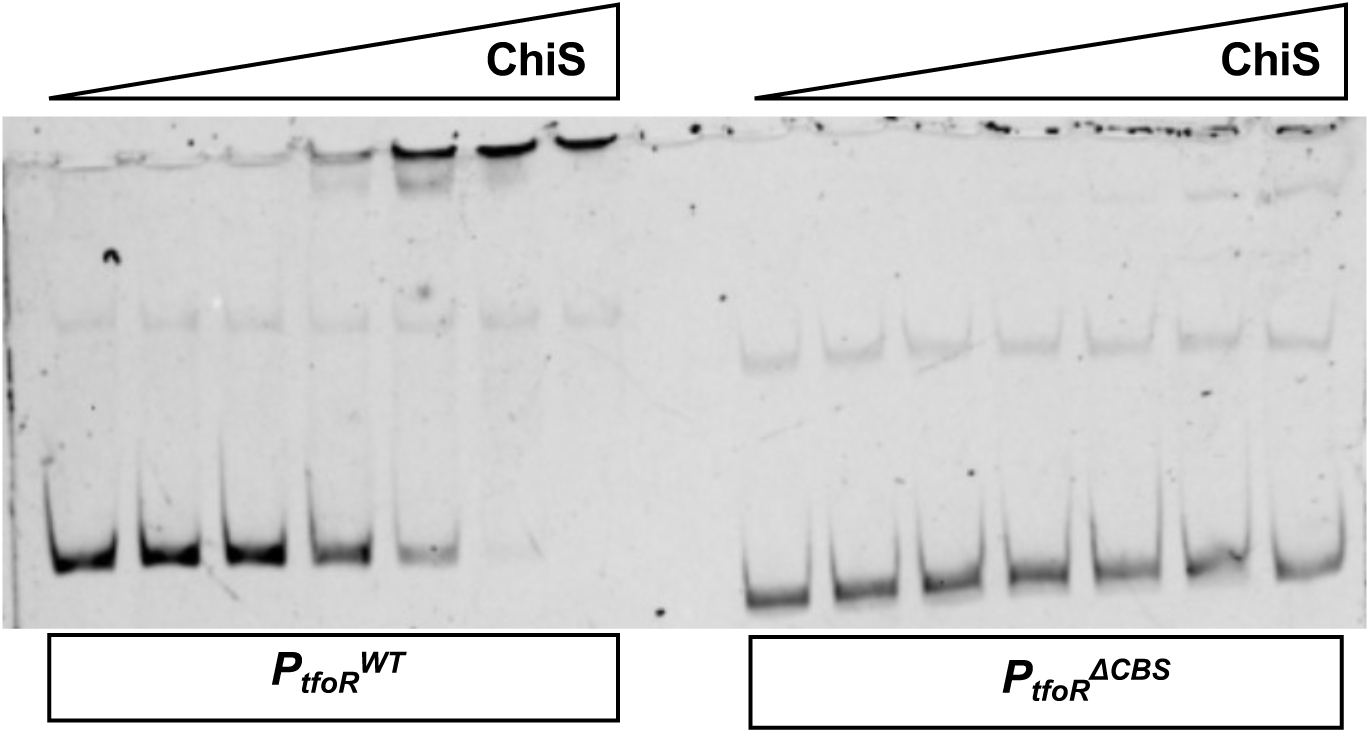
Truncating the 5’ end of P_tfoR_ removes the ChiS binding site. EMSAs of the purified ChiS cytoplasmic domain with the indicated probe DNA. Increasing concentrations of ChiS (from left to right: 0 nM, 25 nM, 50 nM, 100 nM, 200 nM, 400 nM, 800 nM) were incubated with the indicated Cy5-labelled P*_tfoR_* DNA probe. P*_tfoR_*^WT^ denotes the full-length wildtype P*_tfoR_* promoter, while P*tfoR^Δ^*^CBS^ denotes the truncated promoter where the ChiS binding site was removed from P*_tfoR_*. Data are representative of three independent experiments.

**Fig. S7.**
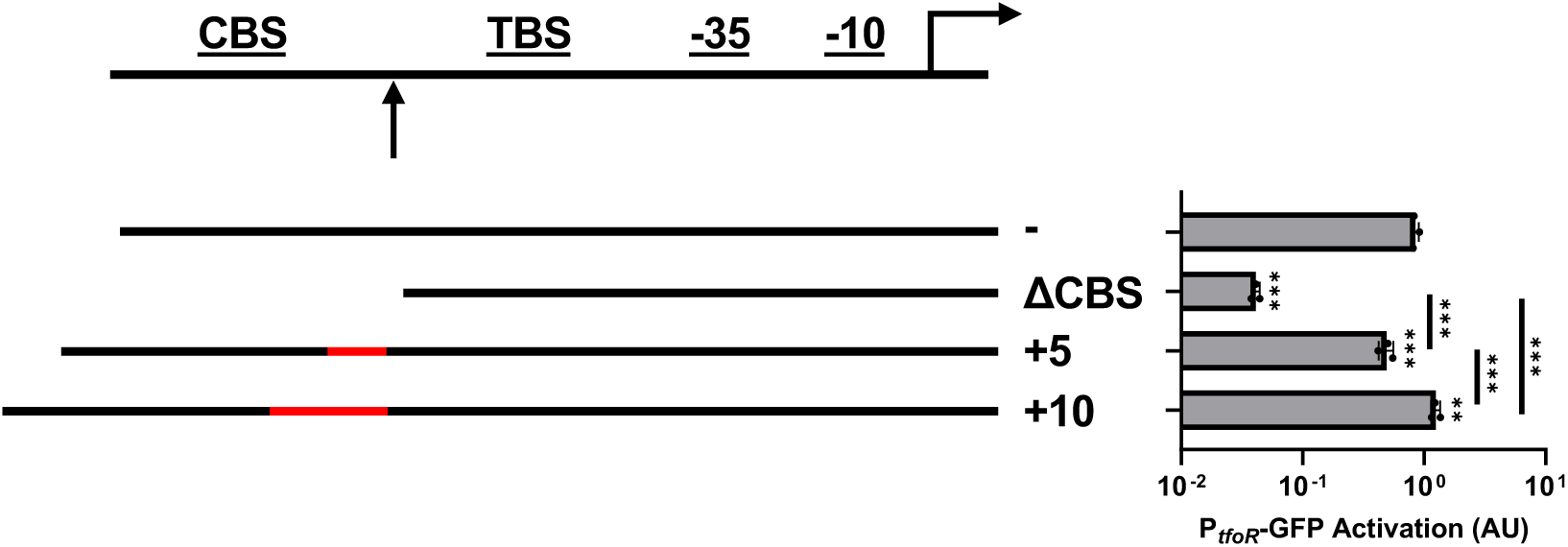
Phasing between the ChiS and TfoS binding sites is not critical for P_tfoR_ activation. Transcriptional reporter assays to assess chitin-dependent gene expression of strains with the indicated PtfoR-*gfp* reporter constructs. Schematic of the P*_tfoR_* reporter constructs depicts where insertions of 5 bp or 10 bp (red) were introduced in between the ChiS binding site (CBS) and TfoS binding site (TBS) (see arrow) to alter the phasing between these elements. Activation of these reporters can be be compared to the parent P*_tfoR_* reporter (positive control) and the P*_tfoR_^ΔCBS^* (negative control). Data are from three biological replicates (300 cells analyzed per replicate) and shown as the geometric mean ± SD. Statistical comparisons were made by one-way ANOVA with Tukey’s multiple comparison test on the log-transformed data (normal distribution confirmed by Shapiro-Wilk test). Statistical identifiers directly above bars represent comparisons made to the parent (top bar). ** = *p* < 0.01, *** = *p* < 0.001.

**Table S1.**
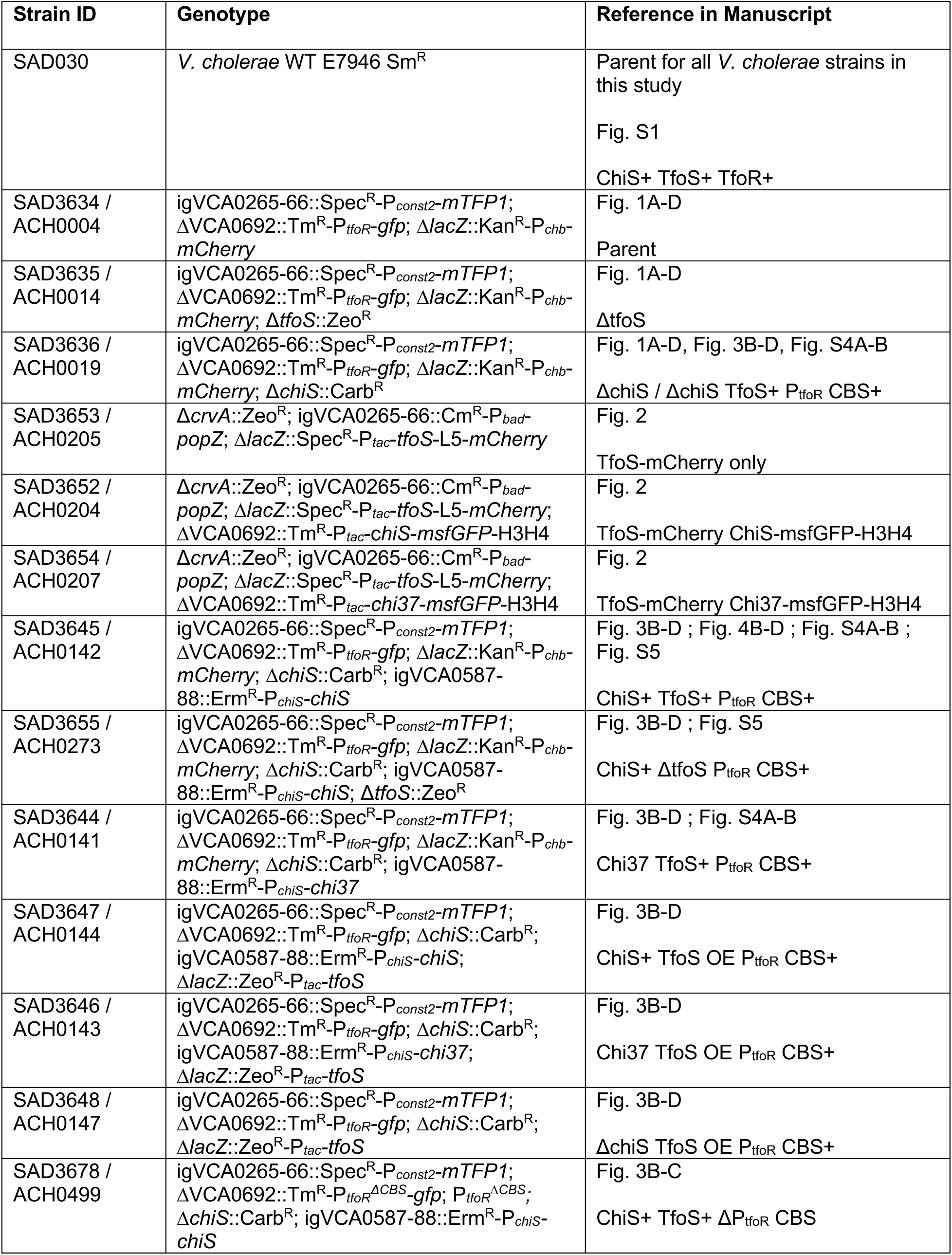
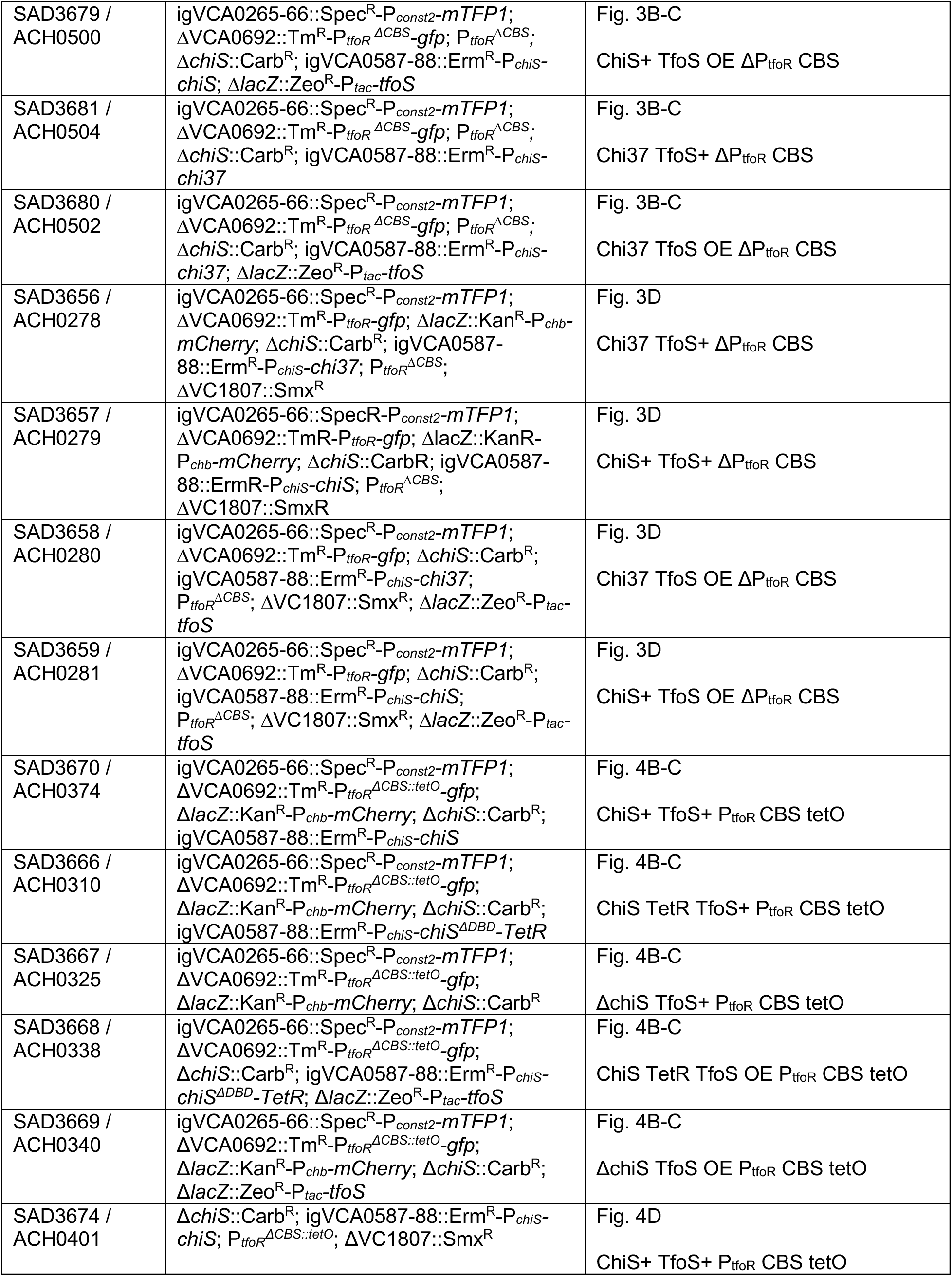
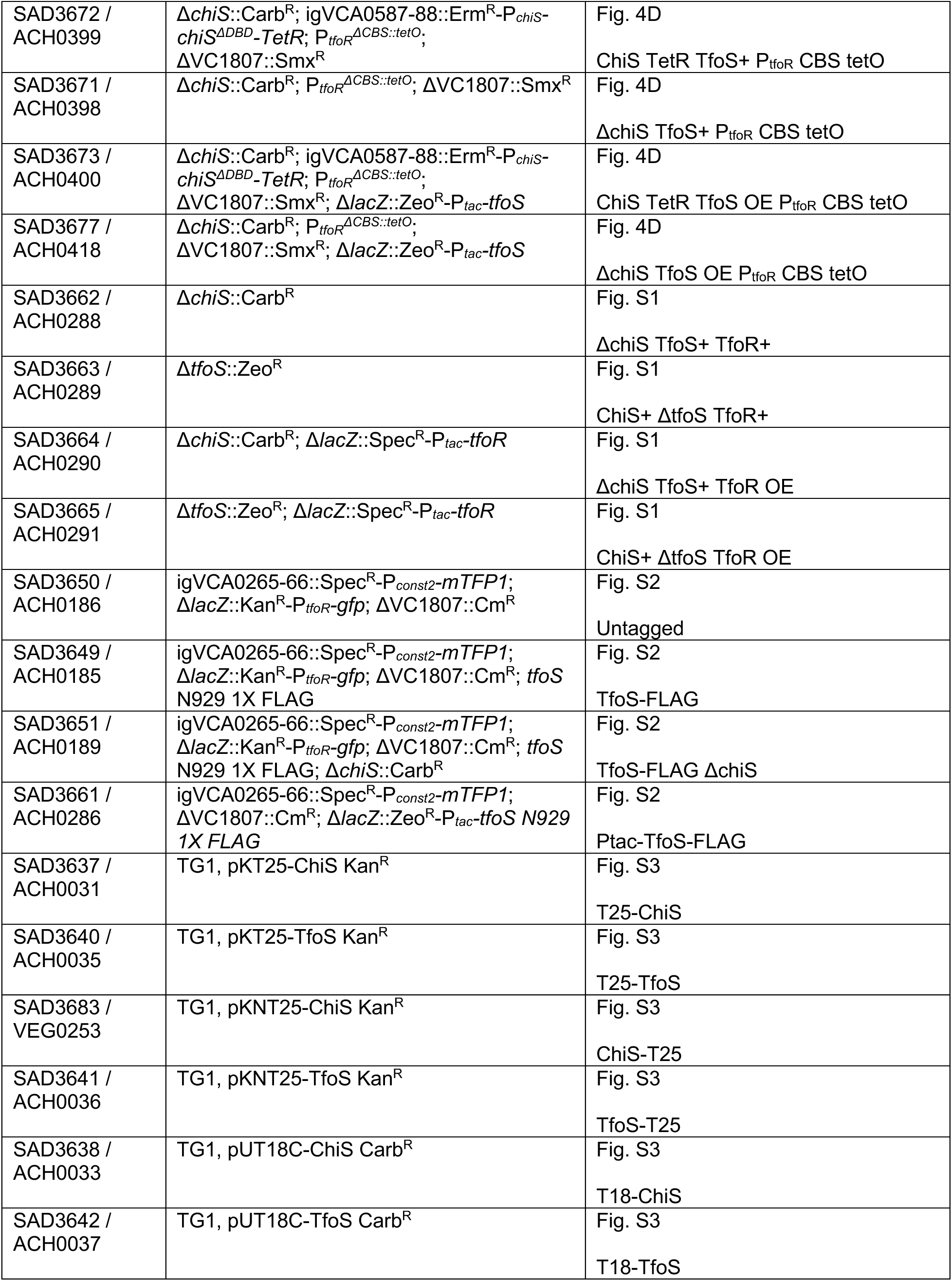
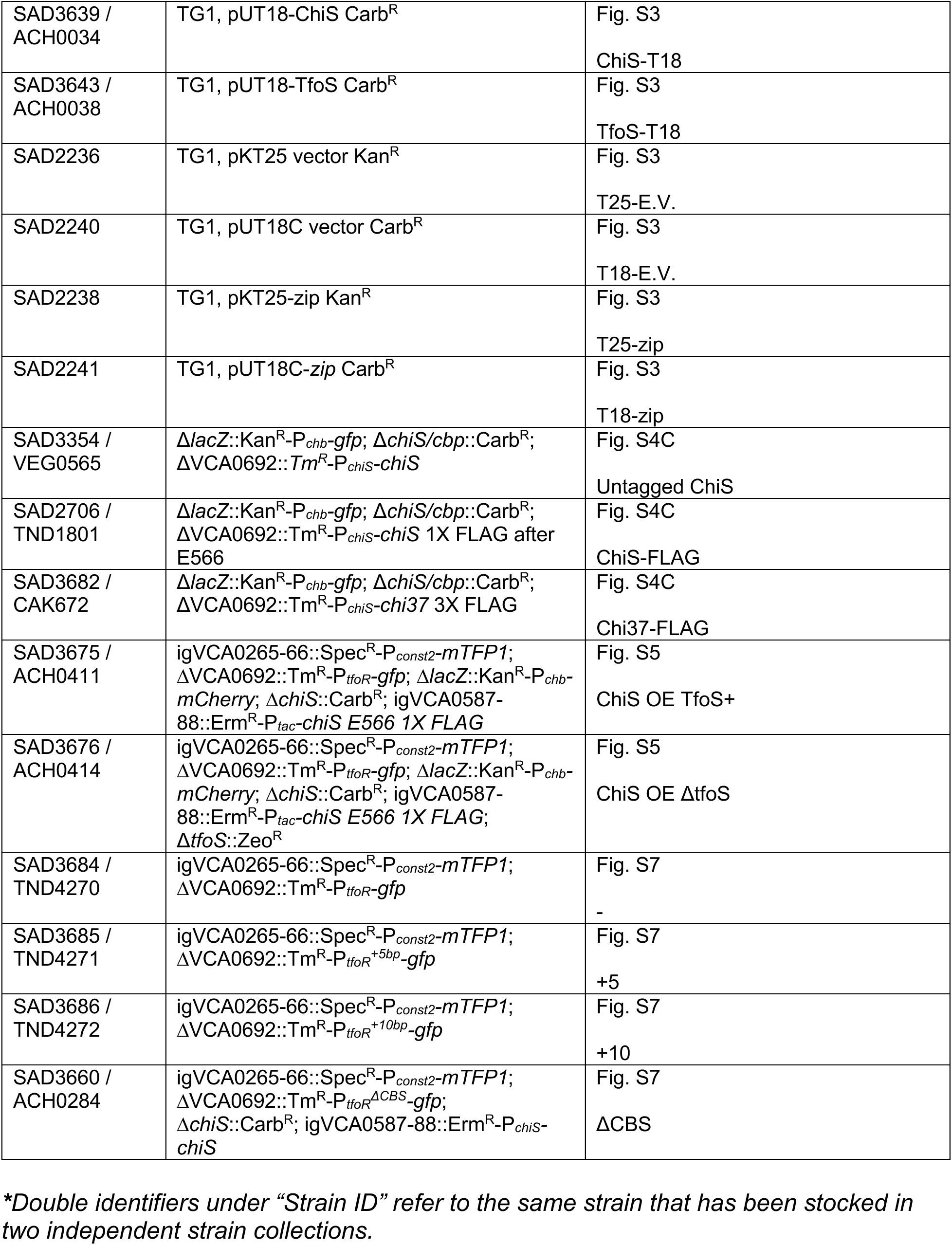
Strains used in this study.

**Table S2.**
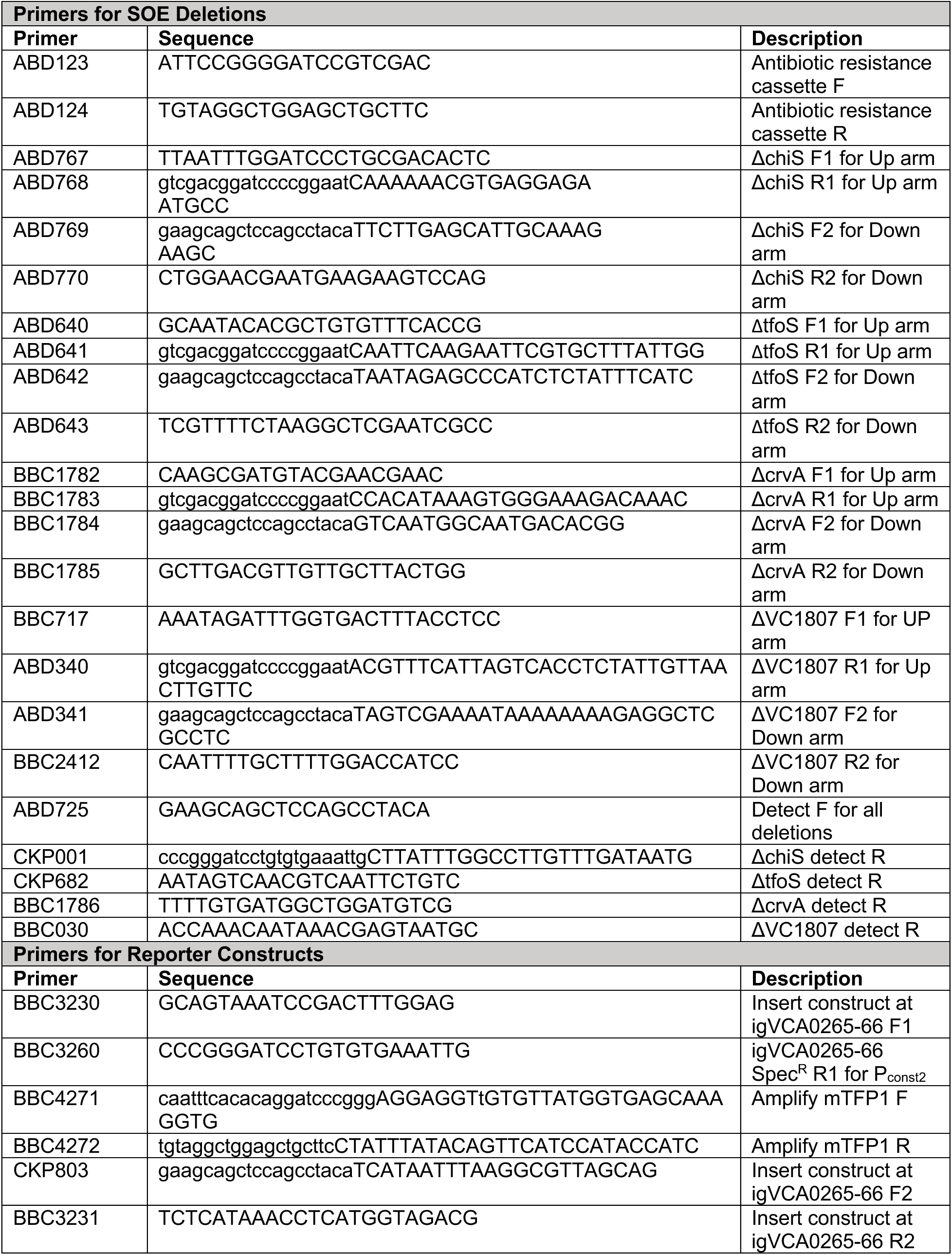
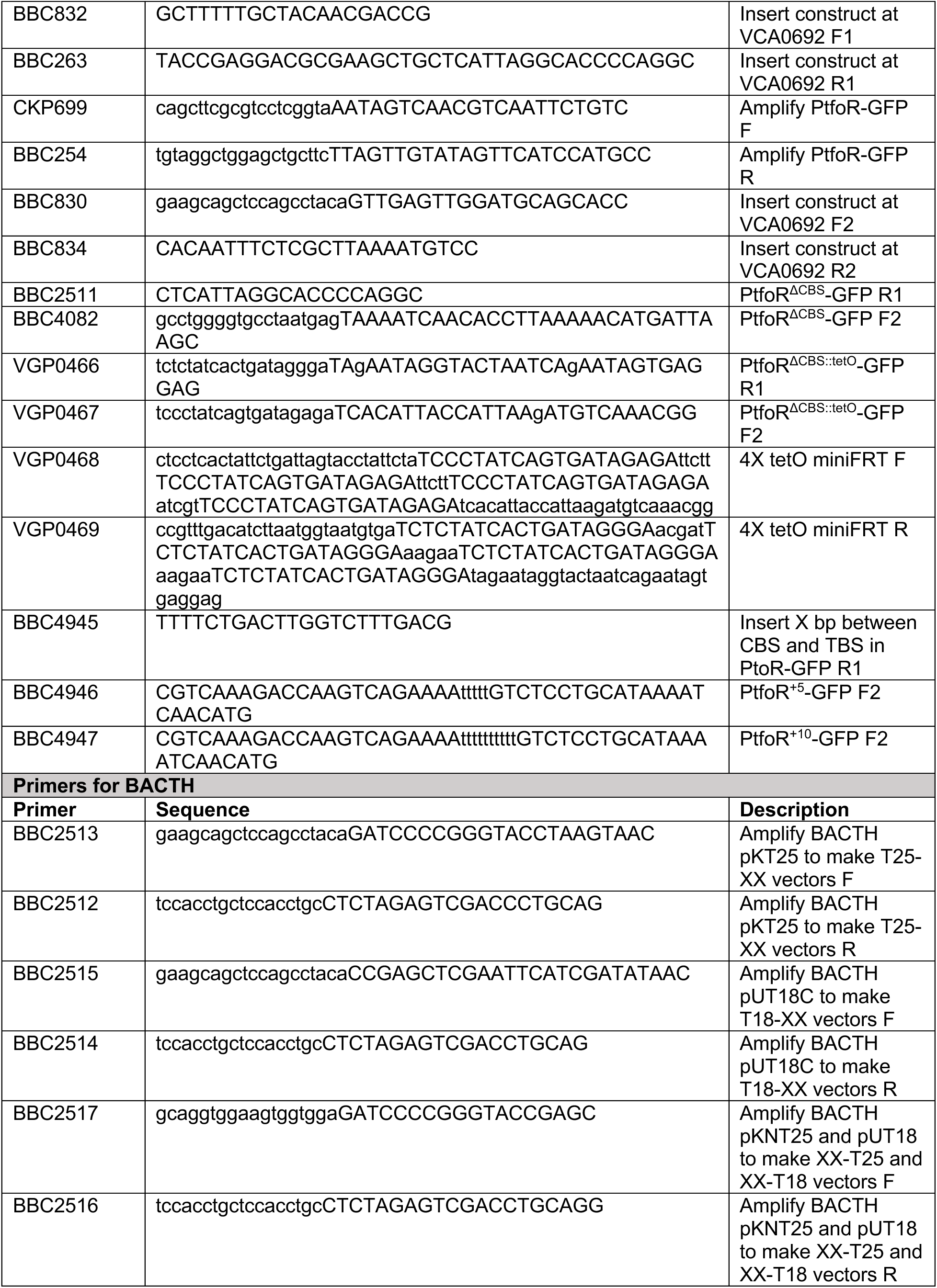
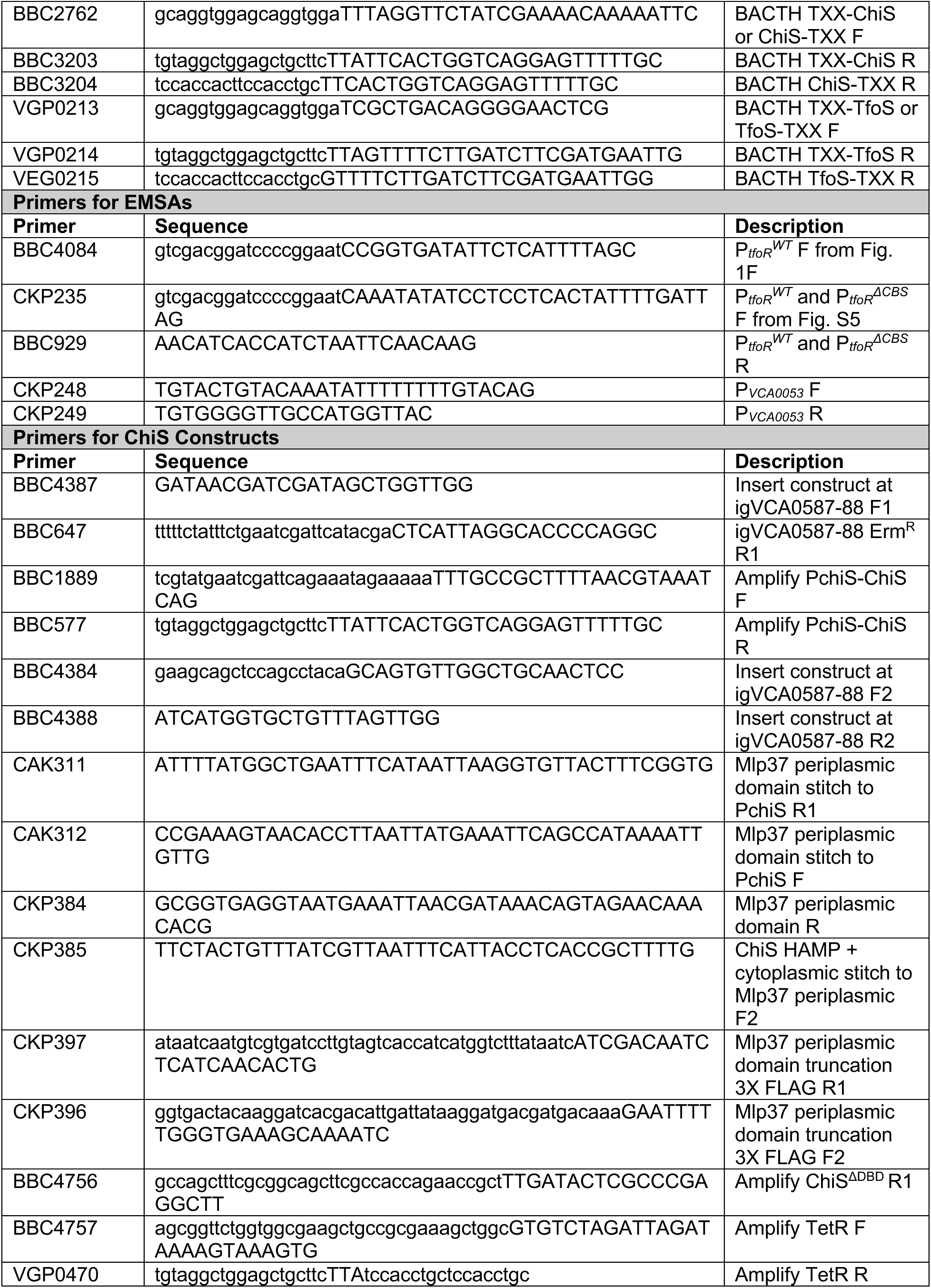
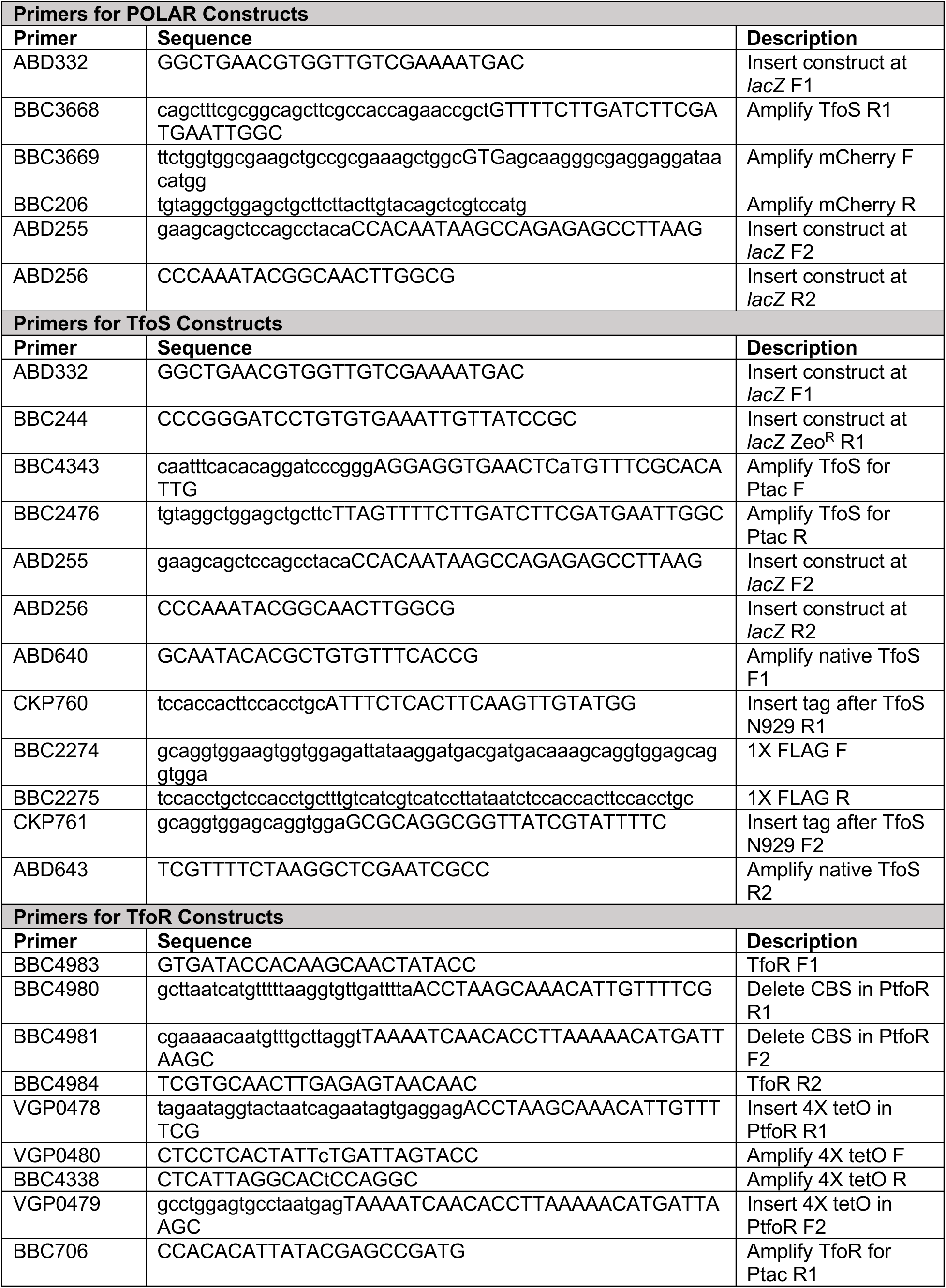
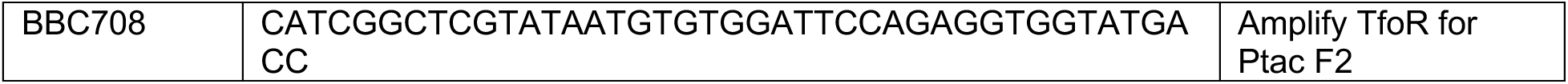
Primers used in this study.

## Notes

### Competing Interest Statement

The authors have declared no competing interest.

### Summary of Updates

Numerous changes to the main text and figures, including the addition of new experiments that further support the main conclusions of the original submission.

